# Colony morphogenesis regulates sporulation dynamics in bacterial biofilms

**DOI:** 10.64898/2026.02.11.705348

**Authors:** Joshua M. Jones, Meiyi Yao, Andrew Mugler, Joseph W. Larkin

## Abstract

In multicellular systems, organized phenotypic heterogeneity emerges from the interplay of processes spanning scales from molecular to population-level. Using *Bacillus subtilis*, we investigated feedback between the collective process of colony expansion and the distribution of spore development among individual cells, a process triggered by starvation. Biofilms are commonly studied using a strain with inhibited sporulation. Intact regulation yielded high-frequency sporulation early in biofilm growth. Biofilm composition was organized by a wave of sporulation driving biofilms toward dormancy from within. However, expansion was also maintained by non-sporulating cells in a narrow front at the external edge. Along with mathematical modeling, we also used mutants with altered biofilm morphogenesis to probe the relationship between colony expansion and sporulation. Sporulation dynamics were patterned by radial expansion, but the faster biofilms spread, the greater the separation of growth and sporulation distributions. We demonstrate essential interplay between cell behavior and the physics of collective expansion that organizes differentiation among cells.

## Introduction

Understanding how genetically identical cells generate and organize phenotypic heterogeneity within groups is fundamental to multicellular biology. Across biological systems, gradients of cues contribute to basic patterning among cells^1^. For bacteria growing in collectives as biofilms, the most abundant mode of microbial growth^2^, self-produced cues are a primary organizer of cell-to-cell phenotypic heterogeneity^3,4^. However, individual cells also contribute to colony-scale processes, such as biomass expansion and morphogenesis. These processes further shape the gradients created and interpreted by cells, which can in turn alter phenotypic patterning^5^. The relative simplicity of biofilm morphology makes their growth a tractable system to study the feedback between cell biology and emergent physics, which together regulate phenotypic composition.

A hallmark of biofilms is the secretion of polymeric substances to form an extracellular matrix^6^. Matrix polymers transform microbial cell collectives into elastic materials whose emergent properties depends on both cell proliferation and matrix composition^7–9^. Biofilm matrix composition is highly studied in *Bacillus subtilis* biofilms. Matrix varies by strain, and biofilms typically contain multiple polymeric substances and other biofilm-specific molecules^10^. Matrix polymers can have global effects on biomass morphology including both promoting and inhibiting radial expansion of colonies depending on the polymer^11,12^. Understanding how cells are reciprocally impacted by the collective process of colony expansion requires real-time mapping of cell states during biofilm growth.

Phenotypic heterogeneity is well-documented in *B. subtilis* biofilms. Cells with distinct metabolic states and specialized functions co-exist, and their abundances vary^13,14^. Biofilms also contain dormant spores^15^, formed in response to high density and starvation. Spore formation in *B. subtilis* is highly studied at the molecular and cellular level. A pathway known as the phosphorelay regulates the activation of sporulation and integrates numerous signals to initiate sporulation when growth becomes less favorable^16,17^. Spore formation is a terminal process for a cell, culminating in death and lysis to release the spore. In biofilms, sporulation is enriched in a distribution consistent with nutrient depletion gradients^13,18,19^. However, measurements of spore prevalence in biofilms come mostly from a single strain of *B. subtilis*, and strains have genetic diversity at various points of sporulation regulation, which could alter timing and patterning.

Here, we use sporulation in biofilms as a model system to probe phenotypic patterning. We show that the distribution of sporulating cells in biofilm colonies is organized by the rate of biomass radial expansion. Central to this observation was comparison of multiple strains of *B. subtilis*. The most studied biofilm-forming strain sporulates late due to an atypical and defective cell-cell signaling regulator. We show that when regulation is intact, sporulation is shifted to the earliest stages of biofilm formation, concurrently with bulk population growth. Sporulation gene expression formed a wave that spread through biomass, rapidly depleting cells from the center of colonies. Despite this, radial expansion was sustained long-term by growing cells limited to a narrow front at the peripheral edge of biofilms, even when the populations were mostly dormant spores. Inspired by this observation, we generated a mathematical model of biofilms as an active fluid whose constituent cells grew and converted to spores at nutrient-dependent rates. We present predictions for biofilm cell-spore composition as a function of radial expansion rate, which we tested experimentally by co-culturing producers and non-producers of a matrix polymer that reduced radial expansion. As expansion rate is increased, sporulation activity spreads through biomass at comparatively slower rates, demonstrating interplay between emergent colony-scale physics and cell biology that flexibly organizes biofilm composition.

## Results

### Cell-cell signaling promotes sporulation early in biofilm growth

Efficient sporulation in *B. subtilis* requires high cell density in addition to nutrient limitation^20^. Cells sense their density by re-importing secreted signal peptides (Phr) that modulate the activity of cognate intracellular response regulator (Rap) proteins^21^ (Figure 1A). Multiple Rap-Phr systems found across strains regulate sporulation^21^. The commonly studied *B. subtilis* strain NCIB3610 (hereafter referred to as 3610) has an atypical Phr-insensitive allele, *rapP^3610^*, that inhibits sporulation even at high cell density^22–25^ (Figure 1B). For this reason, we compared sporulation in biofilms formed by strain 3610 to those formed by a *rapP^3610^* knockout strain (3610Δ*rapP*) and an independent environmental isolate (PS-216) that naturally lacks the *rapP* gene^26,27^.

**Figure 1.**
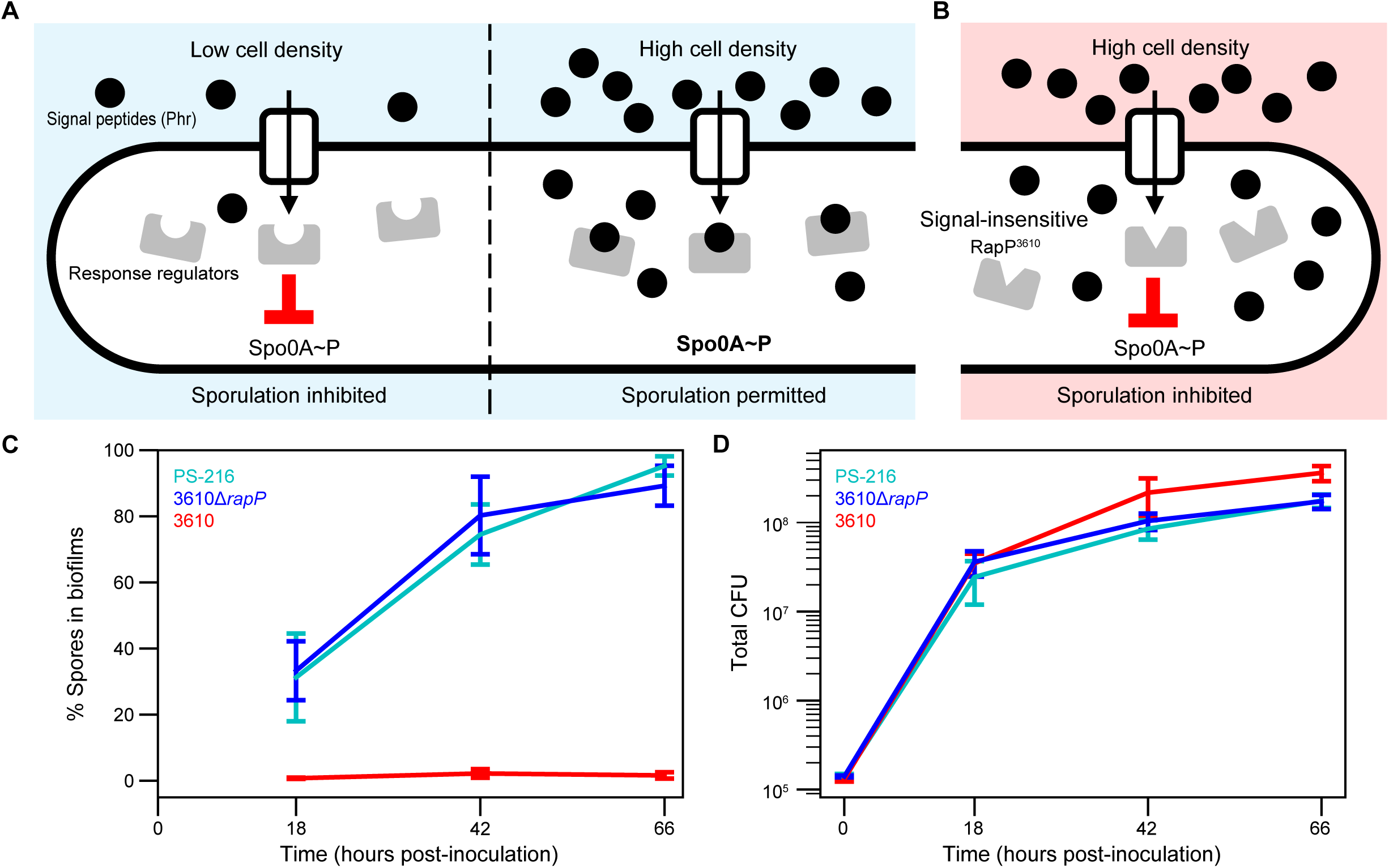
Cell-cell signaling promotes sporulation early in biofilm growth. (A) Schematic of canonical Rap-Phr regulation of sporulation in *B. subtilis*. Intracellular response regulator Rap proteins reduce activation (phosphorylation) of the transcription factor Spo0A, which controls sporulation. Secreted Phr signal peptides accumulate extracellularly in a cell density-dependent manner. Phr peptides are re-imported into cells, where they inhibit the activity of Rap proteins, thereby permitting sporulation at sufficient cell density. (B) Strain 3610 encodes a signal-insensitive Rap protein, RapP, which inhibits sporulation. (C) Percentage of spore-derived CFU and (D) total CFU (cells and spores) in whole biofilms. The central lines represent the mean of 4 replicate biofilms per strain/timepoint (except *n* = 3 for 3610 at 66 hpi). Error bars represent the standard deviation.

We found that the density-sensitive strains (3610Δ*rapP,* PS-216) began forming spores early in biofilm growth. To grow biofilms, we cultured cells on solidified growth medium (MSgg-agarose^15^) at 30℃ (Methods and Materials). We started several replicate populations, which were harvested at intervals to create a time-course. Biofilms were mechanically disrupted and plated to count total viable colony-forming units (CFU) derived from both cells and spores. The percentage of spores in biofilms was determined by measuring heat-resistant CFU as a fraction of total CFU. Spores made up ∼30% of total CFU by 18 hours post-inoculation (hpi) in biofilms formed by 3610Δ*rapP* and PS-216, and by 66 hpi, ∼90% of CFU were spore-derived (Figure 1C). In contrast, biofilms formed by strain 3610 contained less than 2% spores at 66 hpi. Producing a mature spore takes 7-10 hours at 37℃^28–30^. Therefore, in biofilms formed by density-sensitive strains, many cells ceased growth and initiated sporulation as early as 10 hpi. Despite early sporulation, total CFUs increased by a factor of 5-7 between 18 and 66 hpi (Figure 1D). This indicated that a population of growing cells was maintained while many committed to spore development.

### Defective cell-cell signaling alters biofilm morphology

As others have previously reported, we also found that the presence of *rapP^3610^* altered biofilm morphology^23,31^ (Figures 2A and S1A). Strain 3610 biofilms typically increased in radius by over 3 mm within 24 hours, while those formed by 3610Δ*rapP* and PS-216 expanded 4-6-fold less (Figure 2A). They also formed mucoid biofilms in contrast to the ‘dry’ appearance of 3610 (Figure 2A). We verified that *rapP^3610^* was sufficient for both phenotypes (Figure S1). The reported effects of *rapP^3610^* on gene expression are complex; *rapP^3610^* interacts with multiple regulatory pathways that control cell differentiation decisions, including expression of genes for secreted molecules that impact colony expansion^22,23^. The profound impact of *rapP^3610^* on both sporulation and biofilm morphology prompted us to investigate the role of sporulation in radial expansion and to re-examine the roles of secreted biofilm matrix polymers.

**Figure 2.**
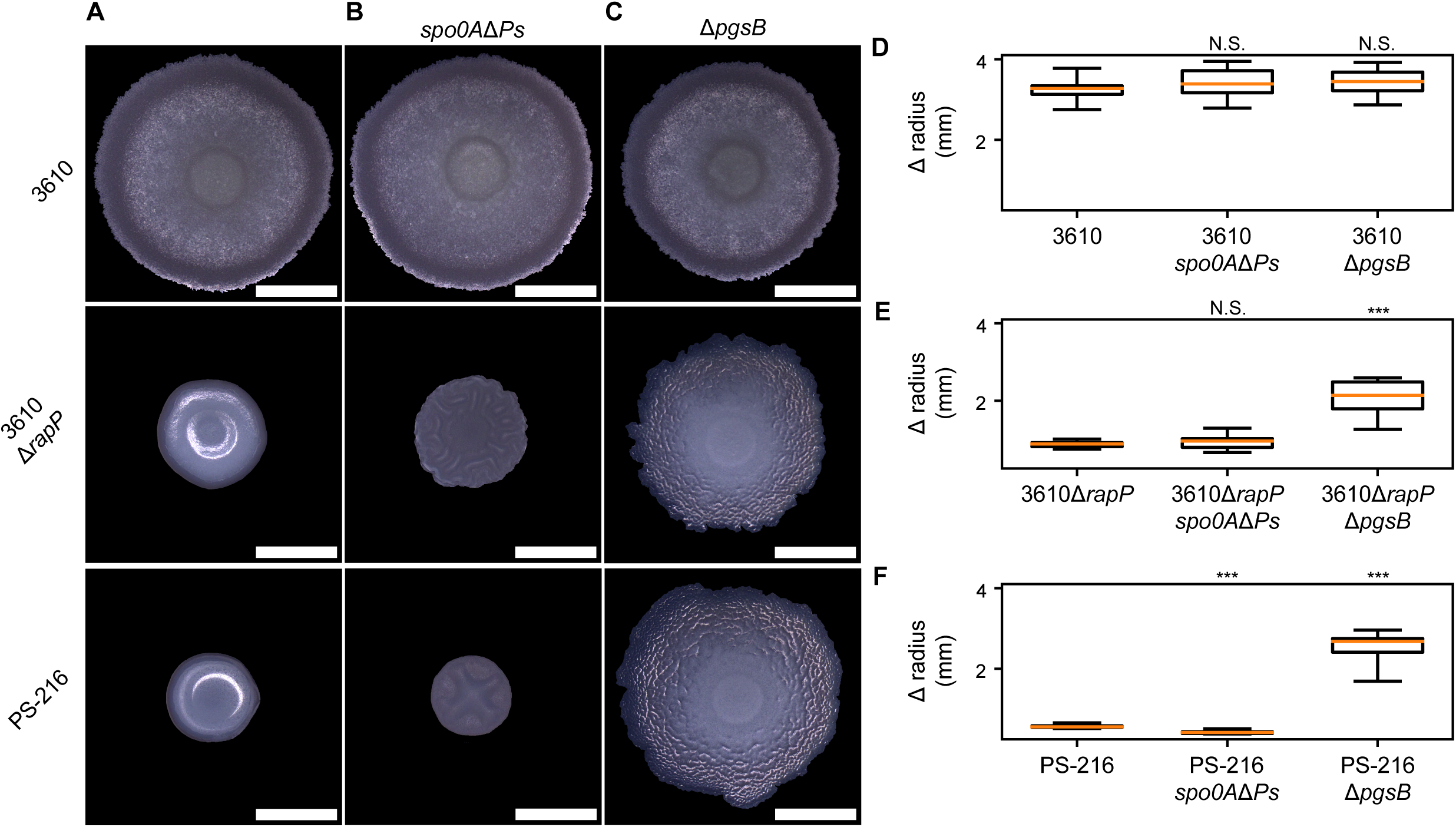
Poly-*γ*-glutamic acid, not sporulation, inhibits radial spreading. (A-C) Color photographs of the top surface of biofilms at 24 hpi. (D-F) Boxplots representing the change in biofilm radius from inoculation to 24 hpi. Central lines indicate medians, boxes enclose the middle 50% of the data, and whiskers indicate the minimum and maximum measurements; *n* = 12 biofilms. *\*\*\*p* < 0.001 by doubled-sided t-test on independent samples with unequal variance (*p* = 0.177 for 3610*spo0A*Δ*Ps*, *p* = 0.152 for 3610Δ*pgsB*, *p* = 0.248 for 3610Δ*rapPspo0AΔPs*, *p* = 8.44 x 10^-7^ for 3610Δ*rapP*Δ*pgsB*, *p* = 8.45 x 10^-9^ for PS-216*spo0A*Δ*Ps*, *p* = 2.56 x 10^-9^ for PS-216Δ*pgsB*). Scale bars 3 mm.

### Sporulation does not inhibit biofilm radial expansion

Cell proliferation generates mechanical force that contributes to physical expansion of biomass. Sporulation in biofilms could decrease radial expansion by depleting the population of actively dividing cells. To test the role of sporulation, we blocked sporulation initiation using a mutation (*spo0A*Δ*Ps*) that reduces the amount of the transcription factor Spo0A, which is required for sporulation^32,33^. The *spo0A*Δ*Ps* mutant was previously shown to form robust biofilms completely lacking spores^34^. In all three strains, blocking sporulation failed to increase biofilm radial expansion (Figure 2B). Since reduction of growth rate triggers sporulation initiation^35,36^, we suspect that sporulation-promoting conditions within biofilms are unsuitable for much additional growth of non-sporulating cells.

### Poly-ɣ-glutamic acid inhibits biofilm radial expansion

Because sporulation had little impact on radial expansion, we investigated the role of matrix polymers. Some strains, but not 3610, produce matrix rich in a hygroscopic polymer, poly-ɣ-glutamic acid (PGA)^37–39^. We suspect that *rapP^3610^* inhibits PGA production for two reasons. PGA was previously linked to mucoid colonies^37^, and we showed *rapP^3610^* eliminated mucoidy (Figure S1A). Second, transient PGA production by strain 3610 early in biofilm growth was shown to temporarily slow radial expansion^12^. Reduced biofilm size and mucoid morphology of 3610Δ*rapP* and PS-216 are both consistent with elevated PGA production in the absence of *rapP^3610^*.

We found that PGA was responsible for restricted radial expansion of biofilms formed by 3610Δ*rapP* and PS-216. To probe the role of PGA, we deleted *pgsB*, which is required for PGA synthesis^40^. Loss of PGA more than doubled radial expansion for 3610Δ*rapP* and PS-216 biofilms at 24 hpi (Figure 2C). As previously reported, Δ*pgsB* had no effect on morphology or total expansion for strain 3610 under these conditions^12,41^ (Figure 2C). We also tested the role of another matrix polymer, extracellular polysaccharide (EPS), which was reported to promote radial expansion of strain 3610 biofilms^11^. We found loss of EPS (Δ*epsA-O*) decreased biofilm expansion only for strains 3610 and 3610Δ*rapP*, and the effect on 3610Δ*rapP* was small (Figure S2). While EPS and sporulation contribute to biofilm morphogenesis for strains without *rapP^3610^*, our measurements show that PGA is the main factor determining biofilm radial expansion (Figure 2DEF).

### Growth and sporulation in biofilms segregate spatially

We showed using direct CFU counts that actively expanding biofilms are composed of a mixture of growing cells and dormant spores. To visualize how sporulation was spatially distributed during biofilm growth, we performed timelapse confocal microscopy using a reporter for sporulation initiation (P*spoIIG-yfp*), which was integrated in the chromosome. A distinct phase of *spoIIG* expression occurs after the end of vegetative growth and commitment to sporulation^42–44^. To generically label cellular biomass, the reporter strain also encoded P*veg-mScarlet*, which is constitutively expressed^45,46^. Biofilm growth was initiated by depositing droplets of suspended cells onto the substrate, which dried as circular patches on the surface. We imaged millimeter-scale regions of interest oriented along the radius of expanding biofilms for up to 48 hours (Figure 3A).

**Figure 3.**
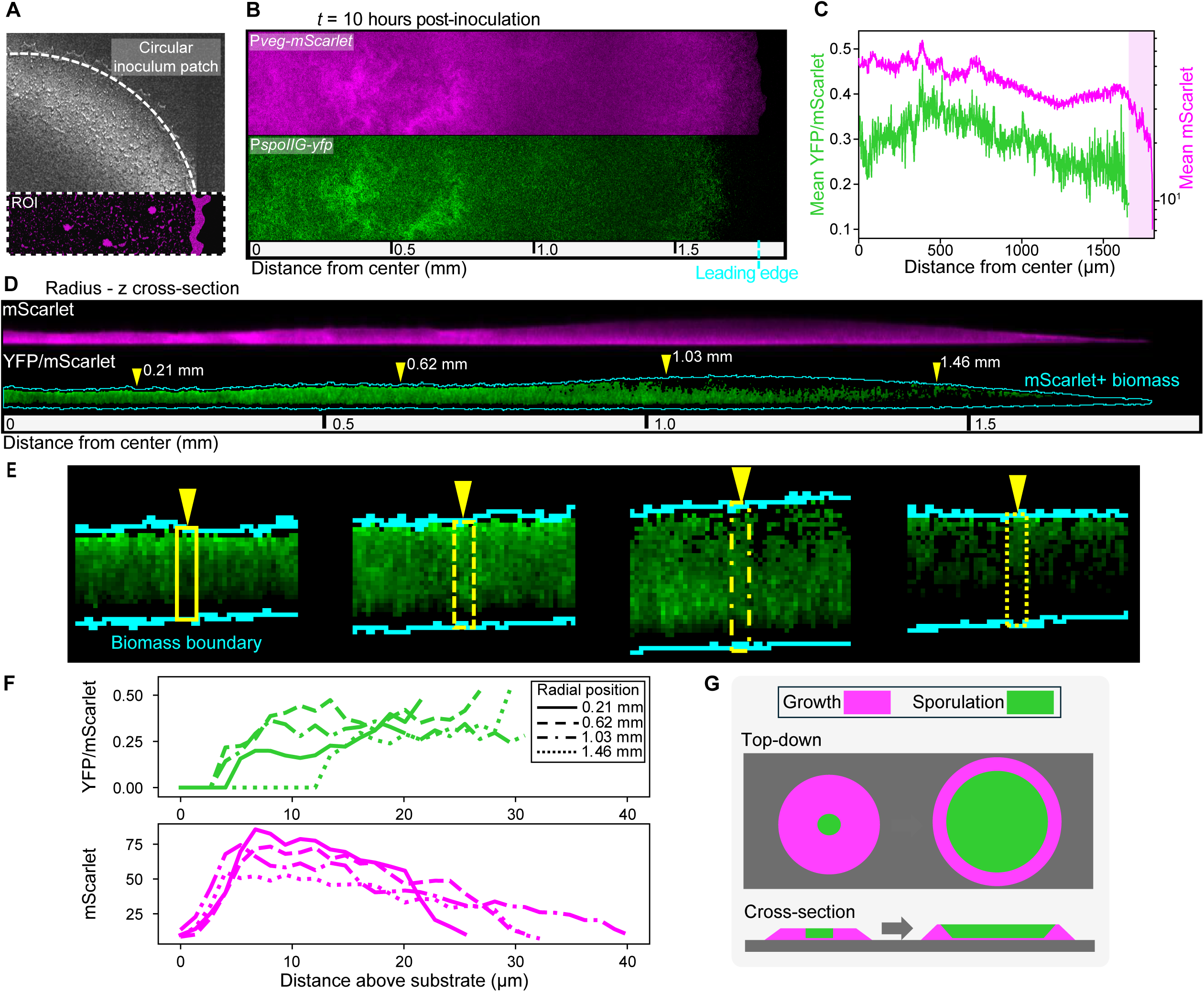
Early spatial separation of growth and sporulation in biofilms. (A) Section of a dried inoculum patch on the substrate surface. The overlayed mScarlet fluorescence data (*t* = 1 hpi) represents a typical ROI oriented along the radius. (B-E) Live confocal imaging of a representative 3610Δ*rapP* biofilm at 25x magnification. (B) Top-down Z-projections made by averaging threshold-passing pixels across z stacks. Magenta, P*veg-mScarlet*; green, P*spoIIG-yfp*. (C) Radial profiles of relative sporulation activity (mean YFP/mScarlet across z stacks for YFP-positive pixels) and P*veg-mScarlet* intensity from a slice (∼40 μm width) through z stacks. The shaded area corresponds to YFP-negative, mScarlet-positive biomass. (D-E) Radius-Z cross-sections made by slice-averaging z stacks. Magenta, P*veg-mScarlet*; green, YFP/mScarlet. The cyan outline surrounds mScarlet-positive pixels. Images in (D) were straightened (ImageJ) to correct for substrate curvature. Yellow triangles specify the positions (distances from biofilm center) of zoom-ins shown in E, which were cropped from original unstraightened data. (E) YFP/mScarlet for YFP-positive pixels. Yellow triangles mark pixel columns measured in F. (F) Intensity profiles of YFP/mScarlet and mScarlet at positions indicated in E. (G) Simplified illustration of the sporulation distribution early in biofilm growth.

We found that by 10 hours of growth, P*spoIIG-yfp* activity had spread from sparse pockets near the biofilm center to the majority of biomass (Figure 3B). To quantify sporulation promoter activity spatially, we measured YFP intensity relative to constitutively expressed mScarlet. This was necessary to compensate for variation in YFP intensity caused by heterogeneous cell density and the effects of growth rate on gene expression and protein accumulation^47,48^. We then averaged YFP/mScarlet values across z stacks to generate radial profiles of relative sporulation activity, which was maximum near the biofilm center and decreased toward the leading edge until reaching a region of cells without detectable sporulation (Figure 3C). We also found that sporulation was enriched in upper layers of biomass. By extracting radius-Z cross-sections from confocal stacks, we found that a small layer of biomass closest to the substrate had little to no detectable P*spoIIG-yfp*, while in upper biomass YFP/mScarlet values saturated (Figure 3D). We quantified this with vertical profiles of pixel intensity spanning from lower to upper boundaries of biomass, as determined by mScarlet thresholding (Figures 3E and 3F). At this early stage of growth, the distribution of sporulation activity in 3610Δ*rapP* biofilms already resembled that of strain 3610 at more advanced stages^13,18^. When disruption of self-sensing is relieved cells sporulate readily, except those with greatest nutrient access along the substrate and at the colony edge (Figure 3G).

### Biofilm expansion is maintained by a front of non-sporulating cells

Despite widespread sporulation beginning early in biofilm growth, vegetative (non-sporulating) cells at the peripheral edge continued to drive biomass spread. Biofilm radius increased by an additional ∼50% (∼0.9 mm) between 10 and 48 hpi. This was led by a propagating front of mScarlet-containing biomass that was sustained as interior biomass lost reporter signal over time (Figure 4A and Video S1). To quantify the evolving distribution of P*veg*-*mScarlet* activity, we summed threshold-passing intensities across z stacks to generate images of total integrated mScarlet. Initially, mScarlet was relatively uniform from the biofilm center to leading edge, but after ∼15 hours the distribution formed a steadily narrowing front at the edge. (Figure 4A and Video S2). Narrowing of the vegetative cell front is consistent with overall nutrient depletion driving increasingly more sporulation at the trailing edge of the front compared to new growth at the leading edge.

**Figure 4.**
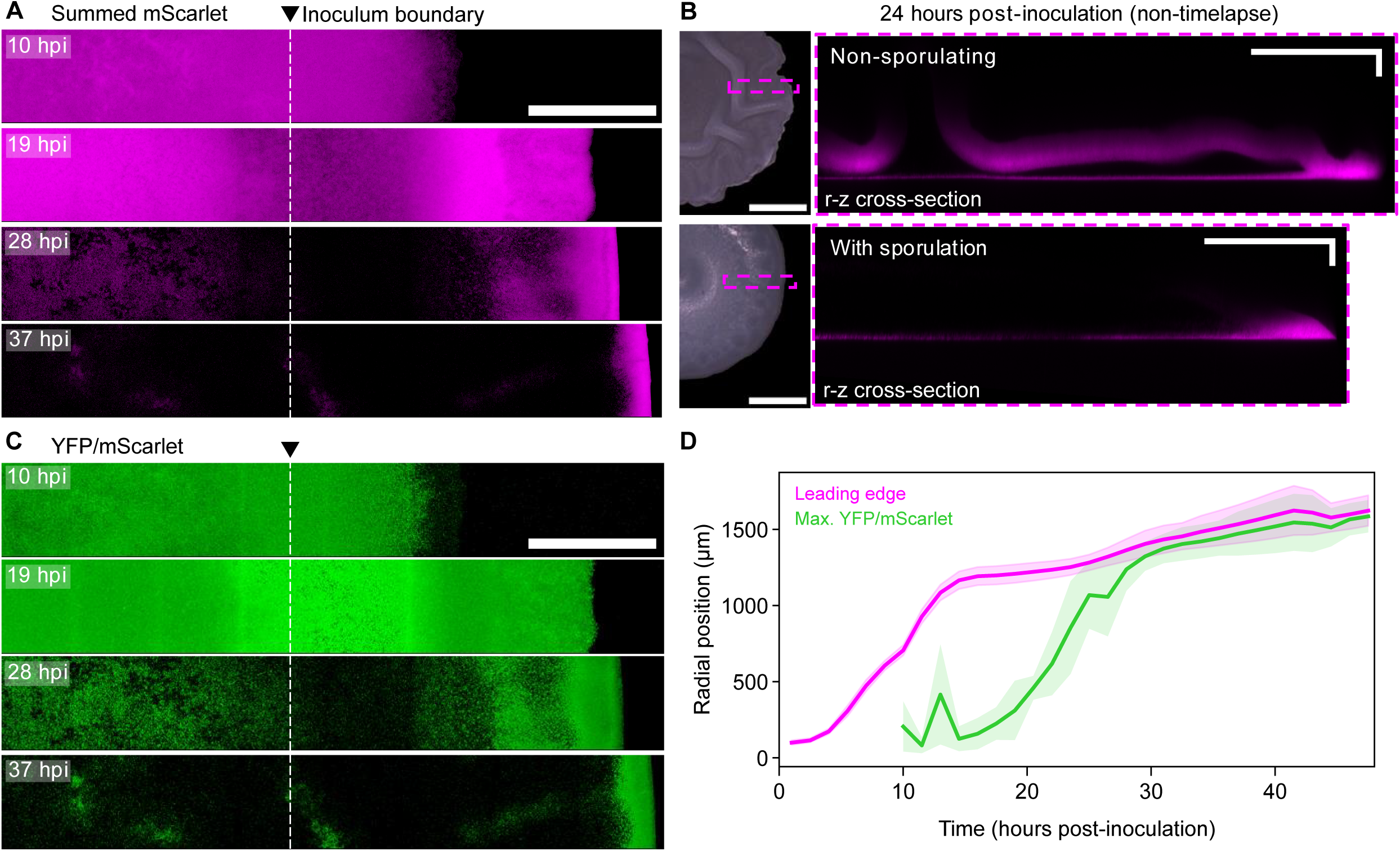
A traveling front of vegetative cells deposits spore-formers. (A) Z-projections generated by summing threshold-passing mScarlet intensities across z stacks for the full time-series dataset corresponding to Figure 3. Images were log-transformed to highlight low intensities. (B) Top-down color photographs (left) and radius-Z cross-sections of mScarlet data (right). Samples were pre-grown for 24 hours prior to imaging. Magenta boxes indicate the areas selected for confocal imaging. (C) Z-projections (corresponding to panel A) generated by averaging YFP/mScarlet across z stacks. YFP/mScarlet was measured for all dual YFP- and mScarlet-positive pixels. Images were log-transformed to highlight small values. (D) Tracking of biomass leading edge position and the position of maximum YFP/mScarlet along biofilm radii. Measurements were taken on 7 radial segments across 4 biofilms. Central lines indicate mean positions, and shading indicates standard deviations. Scale bars (A,C) 0.5 mm; (B) 1 mm (photos), 250 x 50 μm (fluorescence).

We found that loss of P*veg-mScarlet* activity in biofilm interiors was mainly due to sporulation. Biofilms formed by a non-sporulating (*spo0A*Δ*Ps*) version of the reporter strain maintained high mScarlet in interior biomass (Figure 4B, top). Additionally, sporulating biofilms that were pre-grown before imaging (no photobleaching) still had depleted interior mScarlet (Figure 4B, bottom). The *veg* promoter is active during sporulation^45^, and since sporulating cells undergo programmed lysis to release spores, we suspect that mScarlet present in sporulating cells contributes to residual mScarlet in interior biomass. Although we cannot rule out the presence of lingering vegetative cells in the interior of biofilms, our direct CFU counts and timelapse imaging both indicate that most, if not all, biofilm cells eventually sporulate.

### Spread of sporulation has dynamics matching biofilm radial expansion

Within biofilms, cell proliferation drives the formation of nutrient gradients^3,49^. Since nutrient acquisition depends on the size of the circular biofilm ‘footprint’ contacting the substrate in our system, we hypothesized that the dynamics of radial expansion could exert strong feedback on cell behavior, especially sporulation. To investigate this feedback, we quantified the spread of sporulation over time. First, we asked how the initial distribution of cells impacted the sporulation pattern. Inoculated cells were arranged on the substrate as a continuous outer ring with a patchy interior (“coffee stain” pattern, see Figure 3A above) due to evaporation of the inoculum droplet^50^. High levels of sporulation activity first appeared in biomass derived from proliferation and expansion of the outer ring of the inoculum patch (Figure 4C). The role of initial cell density in patterning sporulation was reminiscent of cell death distributions in strain 3610 biofilms determined by initial cell density^51^. After experiencing high YFP/mScarlet, biomass went dark, a reflection of both *spoIIG* downregulation and complete dormancy by the end of spore development. Sporulation and subsequent signal loss spread from the inoculum edge position bi-directionally toward the biofilm center and toward the actively expanding leading edge (Figure 4C and Video S2).

We found that sporulation activity swept through biomass with dynamics that closely resembled those of radial expansion. To compare expansion and sporulation dynamics, we first cropped datasets to examine only the biomass formed by spreading outward from the inoculum patch onto uncolonized substrate. Since YFP/mScarlet profiles generally ascended from low values near the biofilm edge to an interior peak, we quantified spatial spread of sporulation activity by tracking the maximum over time. To mitigate inflation of YFP/mScarlet due to depletion of mScarlet from the interior over time, we excluded measurements beyond the vegetative cell front (Figure S3, Materials and Methods). Despite being spaced apart temporally by ∼12 hours, the position of maximum YFP/mScarlet took a trajectory through biomass that closely matched the past trajectory of the biofilm leading edge (Figure 4D). This inspired a conceptual model of the biofilm edge as a propagating wave of proliferating cells, which at its trailing edge undergoes conversion to spores at a rate dependent on the cell-generated pattern of nutrient depletion. To test this hypothesis and to predict how biofilm expansion relative to evolving nutrient gradients could regulate the patterning of sporulation, we turned to mathematical modeling.

### An active fluid model reproduces linked growth and sporulation dynamics

To model sporulation in growing biofilms, we coupled an active fluid model to the consumption of a diffusing nutrient. Active fluid models have been used extensively to describe biofilm morphological dynamics^11,52–56^. We developed a two-dimensional model, because simple radial propagation sufficed to explain the pattern of sporulation progression. Within simulated biofilms, the rates of both cell growth and their conversion to spores are nutrient-dependent (Figure 5A). Specifically, the growth rate increases with nutrient concentration via a Monod function, and the sporulation rate decreases with nutrient concentration via a Hill function (Figure 5B). Growth leads to biofilm expansion at velocity *v*(*r*, *t*) via a continuity term (Figure 5B), which is the hallmark of an active fluid model. New cellular biomass contributes to radial expansion at a rate *ϕg*, where *ϕ* ≤ 1 is a parameter that allows for tuning how cellular biomass translates to radial expansion. This represents the role of secreted matrix polymers, such as PGA, which alter the rate of radial expansion. Further details of the model, including how the parameter values were estimated and how we solve the model numerically, are provided in the Methods and Materials and Supplemental Information.

**Figure 5.**
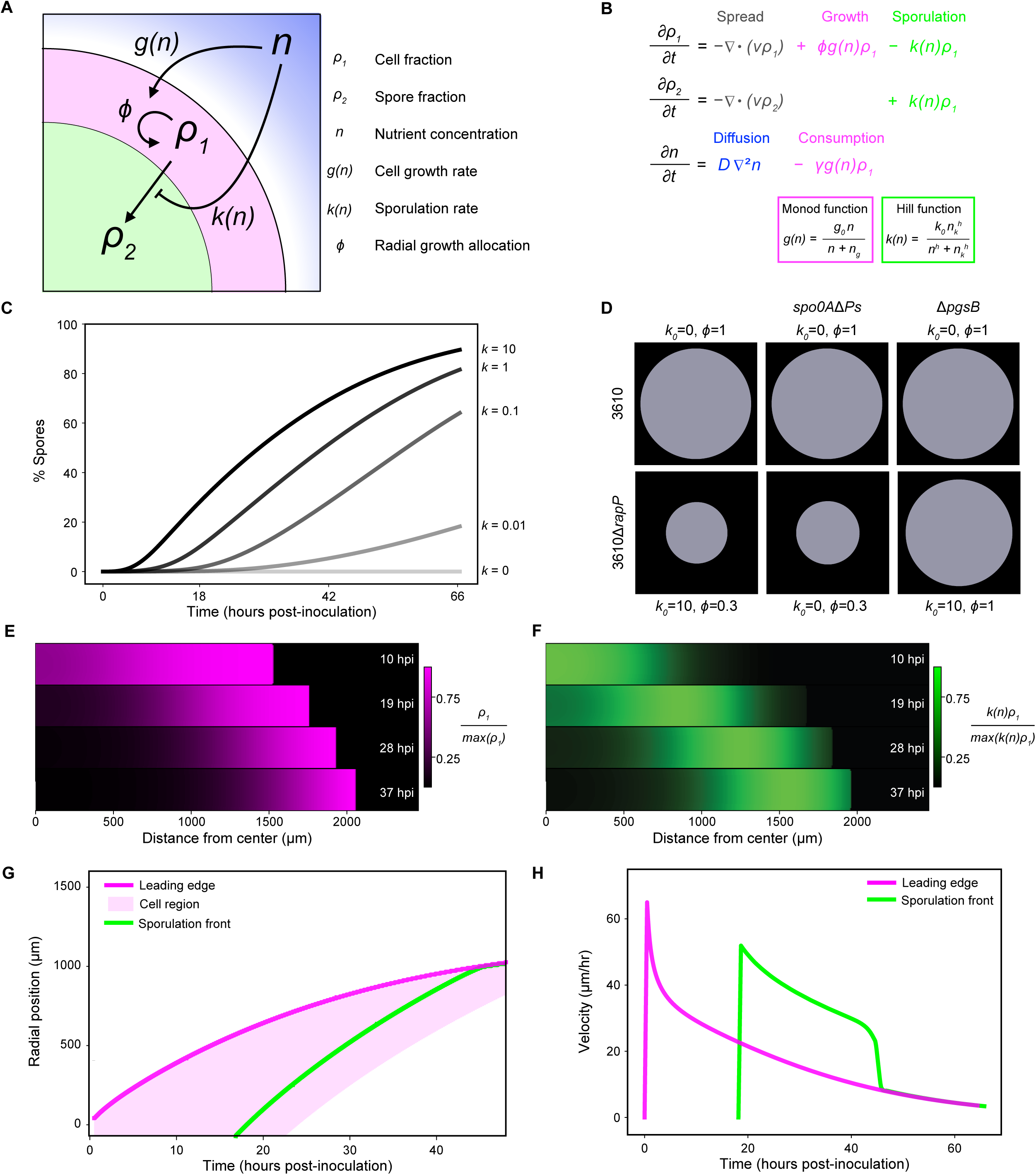
An active fluid model reproduces linked growth and sporulation dynamics. (A) Model diagram. (B) Dynamical equations composing the model. (C) Bulk spore percentage in simulated biofilm populations over time for different *k_0_* values and *ϕ* = 0.3. (D) Representations of total radial expansion (*t* = 66 hpi) for different *ϕ* and *k_0_* values, calibrated to represent the indicated bacterial strains. (E and F) Radial strips of a simulated biofilm (representing 3610Δ*rapP*) at different times. (E) Magenta shows the normalized cell fraction. (F) Green shows the normalized sporulation flux. (G) Change in radial position over time of the leading edge, cell region, and the sporulation front for the biofilm simulated in E and F. The shaded cell region is where the cell fraction of biomass was ≥ 0.5. The sporulation front is defined as the most distant from center position with 75% of the max sporulation flux. (H) Instantaneous velocities (smoothed over a 1-hour window) of the leading edge and sporulation front over time. Initial velocities were set to 0 μm/hr.

We found that the model reproduced the observed feedback between radial expansion rate and sporulation dynamics in biofilms. Tuning *k*_0_, the maximum sporulation rate, allowed us to calibrate the model to represent highly sporulating strains like 3610Δ*rapP* and PS-216 (*k*_0_ = 10) (Figure 5C). Modulating both *ϕ* and *k*_0_ reproduced expansion phenotypes observed in mutant strains, including the minimal impact of sporulation on total expansion (Figure 5D). This occurred because sporulation was offset from growth at the leading edge as in experimental biofilms. Solving the model generated spatial profiles that were directly comparable to experimentally measured fluorescent reporters. Specifically, *ρ*_1_ corresponds to the constitutive (P*veg*) reporter, and the sporulation flux, *kρ*_1_, corresponds to the sporulation initiation (P*spoIIG*) reporter. We found that the model recapitulated the formation of a propagating front of cells that narrowed over time (Figure 5E) and the corresponding broad wave of sporulation flux underlying that cell depletion (Figure 5F). Consistent with experimental biofilms, the sporulation flux wave followed the trajectory of the biofilm leading edge after a time delay (Figure 5G and 5H).

### Model predicts relatively slower spread of sporulation with faster expansion

To predict how the relationship between biomass expansion and sporulation dynamics is affected by the expansion rate itself, we altered radial expansion in the model with the parameter *ϕ* (while maintaining the same intrinsic sporulation rate) and tracked waves of sporulation flux. Increasing *ϕ* resulted in greater average biofilm expansion velocity as well as a larger range of expansion velocities (Figure 6A). Specifically, at large *ϕ*, radial expansion started fast and then slowed, whereas at small *ϕ*, radial expansion began slow and slowed further (Figure 6B, magenta curves). Across values of *ϕ*, sporulation flux propagated toward the leading edge with dynamics matching biofilm radial expansion dynamics, however, with greater *ϕ* (faster expansion) the more closely they agreed (Figure 6B). To quantify this, we discretized radial positions of simulated biomass, and for given positions we compared the instantaneous velocities of the sporulation flux wave and radial expansion.

**Figure 6.**
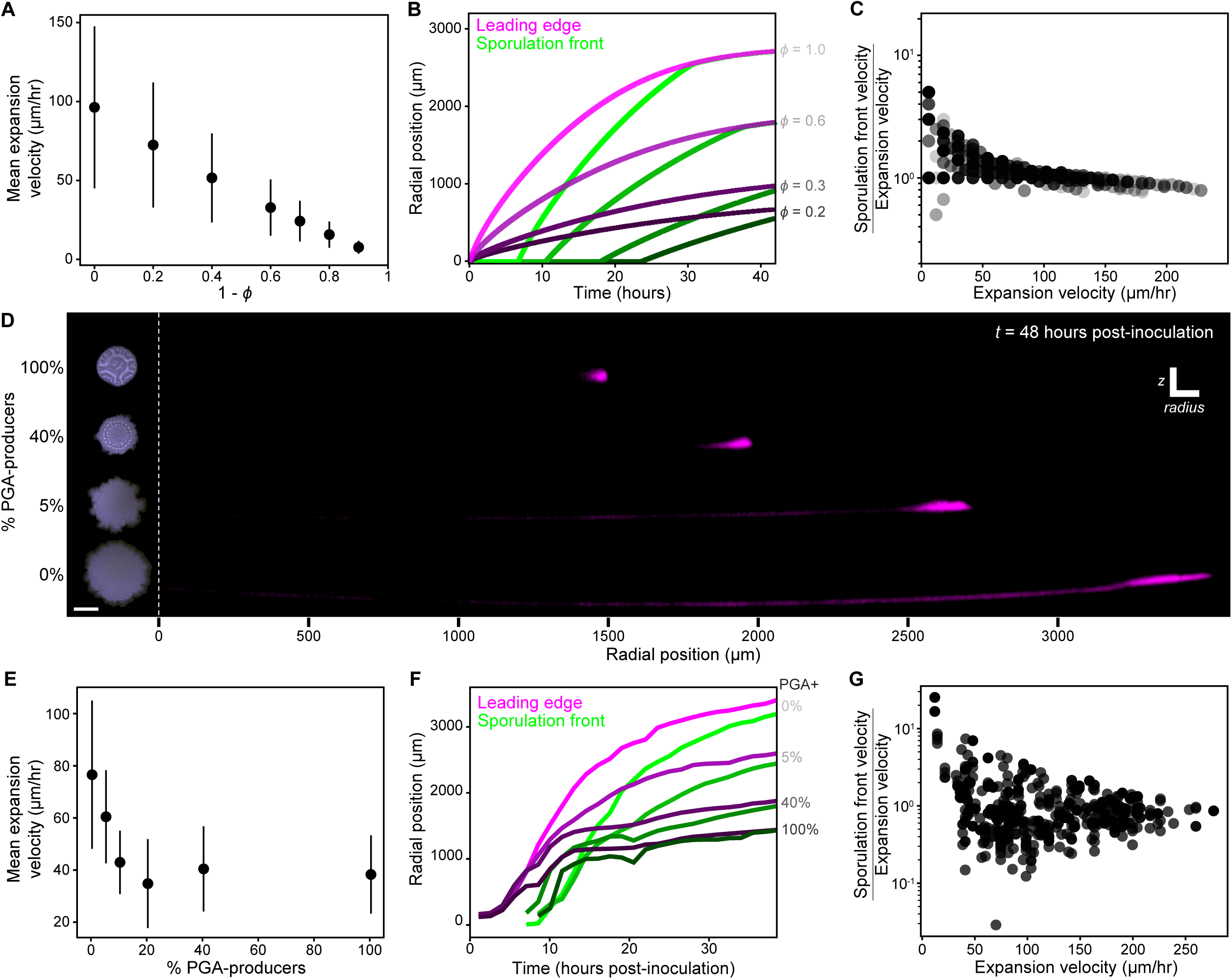
Tuning radial expansion with strain co-culture regulates sporulation dynamics. (A) Biofilm radial expansion velocities (mean ± standard deviation) from simulations with the indicated *ϕ* values. Velocities are calculated from the change in radius (Δ*R*) of the leading edge within a time window of 1 hour. (B) Change in position of the leading edge and the sporulation front for biofilms with the indicated *ϕ* values. Shading levels denote paired curves, with darker shading for lower *ϕ* values. (C) Calculated velocity ratios (sporulation front over expansion) plotted against the corresponding expansion velocities (using instantaneous velocities pooled from the 7 simulated *ϕ* values represented in B). (D) Mixed culture of PGA-producers (3610Δ*rapP*) and non-producers (3610Δ*rapP ΔpgsB*). (Left) Color photographs of the top surface of representative biofilms. (Right) Radius-Z cross-sections of P*veg-mScarlet* for radial segments of biofilms. Images were straightened (ImageJ) to correct for substrate curvature. Radial position is noted with reference to the initial leading edge at *t* = 1 hpi. (E) Radial expansion velocities (mean ± standard deviation) were derived from total radial expansion measurements of 44 biofilms across 2 experiments (0% PGA+, *n* = 5; 5% PGA+, *n* = 7; 10% PGA+, *n* = 8; 20% PGA+, *n* = 8; 40% PGA+, *n* = 8; 100% PGA+, *n* = 8). (F) Change in position of the leading edge and sporulation front for representative biofilms in D. Shading levels denote paired curves, with darker shading indicating greater percentage PGA-producer. (G) Calculated velocity ratios (sporulation front over expansion) plotted against the corresponding expansion velocities using instantaneous velocity data pooled from 20 mixed and mono-culture biofilms from one experiment (0%PGA+, *n* = 1; 5%PGA+, *n* = 3; 10%PGA+, *n* = 4; 20%PGA+, *n* = 4; 40%PGA+, *n* = 4; 100%PGA+, *n* = 4). Scale bar 3 mm (D).

Sporulation flux waves were slowed relative to the radial expansion rate as radial expansion was increased. For large expansion velocities, the ratio of sporulation wave velocity to expansion velocity was near one. In contrast, for small expansion velocities, this ratio increased up to ∼5 (Figure 6C). The non-linearity of this relationship demonstrates the predicted interplay of expansion and sporulation. With slow expansion, propagating sporulation flux waves gained on the leading edge. With faster expansion, the rates converged: the cell front at the edge did not completely outpace sporulation flux waves, which also increased speed but to a lesser extent. Thus, radial expansion rate is predicted to systematically alter the distributions of growing vs sporulating cells.

### Tuning radial expansion with strain co-culture regulates sporulation dynamics

To experimentally probe the predicted feedback between biofilm radial expansion and sporulation dynamics, we took advantage of the rapid spreading phenotype of Δ*pgsB* (PGA-deficient) mutants. Because PGA is secreted into the extracellular environment, we hypothesized that PGA non-producers would still be affected by its presence. Consistent with that hypothesis, we found that mixed culture of PGA-producers and non-producers resulted in intermediate biofilm radial expansion. We made a Δ*pgsB* derivative of our dual *veg*/*spoIIG* reporter strain and prepared mixed inoculums in varying proportions with the original (PGA-producing) reporter strain. Increasing the initial PGA-producer fraction resulted in smaller biofilms (Figure 6D left). Even a small proportion of PGA-producers significantly reduced mean expansion velocity (Figure 6E), allowing us to tune through a broad range of expansion dynamics. We then performed timelapse imaging of sporulation activity in biofilms as described above.

Tuning radial expansion with mixed culture yielded multiple points of agreement with the model. In particular, the relationship between expansion velocity and later propagation of sporulation activity was consistent with the model’s predictions. As before, we measured changes in the distribution of sporulation activity (YFP/mScarlet) during growth. Since lower PGA biofilms maintained mScarlet intensity longer in interior biomass (Figure 6D right and Video S3), YFP/mScarlet profiles tended to form broad plateaus (instead of peaks). As a result, simply tracking the maximum (as in Figure 4D) was noisy and poorly represented the steady spread of high sporulation activity toward the leading edge. Instead, we defined and tracked a ‘sporulation front’ that reliably marked the transition point to high sporulation interior biomass (Methods and Materials). Across biofilms with distinct radial expansion dynamics, the trajectory of the sporulation front roughly matched that of the leading edge (Figure 6F). To directly compare trajectories, which were offset in time, we mapped the instantaneous velocities of radial expansion and the sporulation front to their radial positions (Methods and Materials, Figure S4). Consistent with the model, we found that biomass formed via fast radial expansion experienced sporulation wave propagation at rates close to or less than the expansion velocity (Figure 6G). In contrast, slow expansion velocities (mainly from higher PGA biofilms) were linked to relatively higher sporulation wave velocity. Simply put, regardless of ability to spread as a collective, sporulation inevitably swept through biofilms. Despite that, the collective spreading behavior itself feeds back onto the dynamic distributions of cell phenotypes.

## Discussion

Using an experimental system of radially expanding biofilm colonies, paired with an active fluid mathematical model, we showed that cells differentiated into spores according to a spatial-temporal pattern established by the rate of radial expansion. Further, we showed how cells changed the colony expansion rate with secreted matrix polymers, which in turn altered how sporulation activity was distributed in biofilms. The agreement between our simplified three-component (nutrient, cells, spores) model and experiments manipulating radial expansion argues that self-generated nutrient gradients play a dominant role in patterning phenotypic differentiation.

Essential to our study was the comparison of multiple *B. subtilis* strains, which revealed how a defect in self-sensing due to a gene (*rapP^3610^*) present in a commonly studied strain nearly eliminated sporulation under conditions where a natural environmental isolate without *rapP^3610^* sporulated highly and rapidly. This supports the view that self-sensing, especially with dedicated cell-cell signaling cues, is central to bacterial multicellular behavior^57^. When cells are able to self-sense, sporulation was an inevitable development trajectory for biofilm-resident cells, contrasting with views of sporulation as a stochastic outcome^58^ or a late event of biofilm maturation^13^. Delaying sporulation provides a selective advantage, as long as conditions support cell growth and viability^34,35^. Consistent with that, others have shown that strain 3610 outcompetes natural isolates in co-culture^31^. However, in environments without stable growth conditions, coordinating sporulation such that populations always contain spores regardless of total size could be vital to ensure survival.

Comparison of multiple strains also revealed how *rapP^3610^* alters the colony-scale morphology of biofilms. This was likely due in large part to inhibition of PGA production, which is common in natural isolates lacking the *rapP^3610^* allele^37–39^. We showed that increasing radial expansion rate by reducing PGA content had feedback on phenotypic composition, increasing the spatial-temporal separation between growth and sporulation as cells at the leading edge better accessed nutrients. This suggests suppression of PGA production provides a secondary growth benefit via spreading that could partially explain selection for the *rapP^3610^* variant, independent of its direct inhibitory effect on the sporulation pathway^22^. Consistent with the benefits of radial spreading, experimental lab domestication of a natural *B. subtilis* isolate yielded faster expanding colonies, which was linked to mutation of *degU*^59^. DegU regulates a suite of genes including the *pgsBCAE* genes^60^, which specify PGA production and secretion. Though PGA-producers might sacrifice growth potential in colonies due to impeded expansion, PGA enhances *B. subtilis* plant root colonization^61^ and contributes to desiccation resistance^12^, both essential in soil environments where *B. subtilis* is commonly found. PGA is produced by other organisms (mostly *Bacillus* spp.), both as a secreted polymer and as a capsule (as for *B. anthracis*) that contributes to virulence^62^. The likely prevalence of biofilm forming bacteria that produce both PGA and large numbers of spores calls for investigation into new spore dispersal mechanisms that consider the distinct properties of PGA-rich biomass.

Feedback of physical biomass expansion onto the distributions of cellular phenotypes demonstrates the essential interplay of cell biology and emergent physics in organizing multicellular systems. Others have also observed how radially propagating signals organize phenotypes among cells in radially growing colonies^18,63–65^, arguing that proliferation-driven spreading and the resulting nutrient depletion profile is a fundamental source of pattern generation in biofilms. Adding to this, we also show how other cell activities, specifically matrix secretion, can stretch or compress phenotypic distributions by altering colony-scale expansion. Additional levels of interaction and feedback from other emergent aspects of biofilms, including mechanical gradients^66^ and metabolic cooperation^67,68^ are inevitable. As highlighted by our work, bridging disparate biological and physical scales is needed to understand feedback loops that define the composition and emergent functions of microbial collectives.

## Methods and Materials

### Media and growth conditions

Prior to experiments, cells were grown as light lawns for approximately 24 hours at room temperature on 1.5% agar plates containing 1% w/v glucose, 0.1% w/v monosodium glutamate, and 1x Spizizen’s salts (2 g/l (NH_4_)SO_4_, 14 g/l K_2_HPO_4_, 6 g/l KH_2_PO_4_, 1 g/l Na_3_citrate-2H_2_O, and 0.2 g/l MgSO_4_-7H_2_O)^69^. Light lawns were used to bypass liquid culture steps commonly used to prepare cells, as this selects for biofilm-deficient mutants.

Biofilms were grown at 30°C on 1.5% agarose containing MSgg medium (as defined in Branda et al., 2001 except for tryptophan and phenylalanine, which we did not include). MSgg-agarose was deposited (0.4 ml per well) into glass-bottomed black-wall 24-well plates (Falcon), which were pre-warmed at 70°C to prevent immediate cooling of the agarose. Plates were left open at room temperature for 25 minutes to allow for solidification and drying of the media. Media were prepared immediately before inoculation for each experiment. When indicated, Isopropyl β-d-1-thiogalactopyranoside (IPTG) was added at a final concentration of 10 μM.

Antibiotics were used at the following concentrations for selection on LB-agar plates: chloramphenicol (5 μg/ml), kanamycin (5 μg/ml), spectinomycin (100 μg/ml), tetracycline (12.5 μg/ml), and a combination of erythromycin (0.5 μg/ml) and lincomycin (12.5 μg/ml) to select for macrolide-lincosamide-streptogramin (mls) resistance.

### Strains and alleles

The *B. subtilis* strains used are listed in Table 1. Standard techniques were used for cloning and strain construction^69^. We used a transformable derivative of strain NCIB3610^70^, which was obtained from the Bacillus Genetic Stock Center (BGSC). Some alleles were previously described and are summarized below. The *spo0A*Δ*Ps* allele was derived from AG1242^32^. The P*spoIIG-yfp* reporter was derived from AES3140^71^.

**Table 1.**
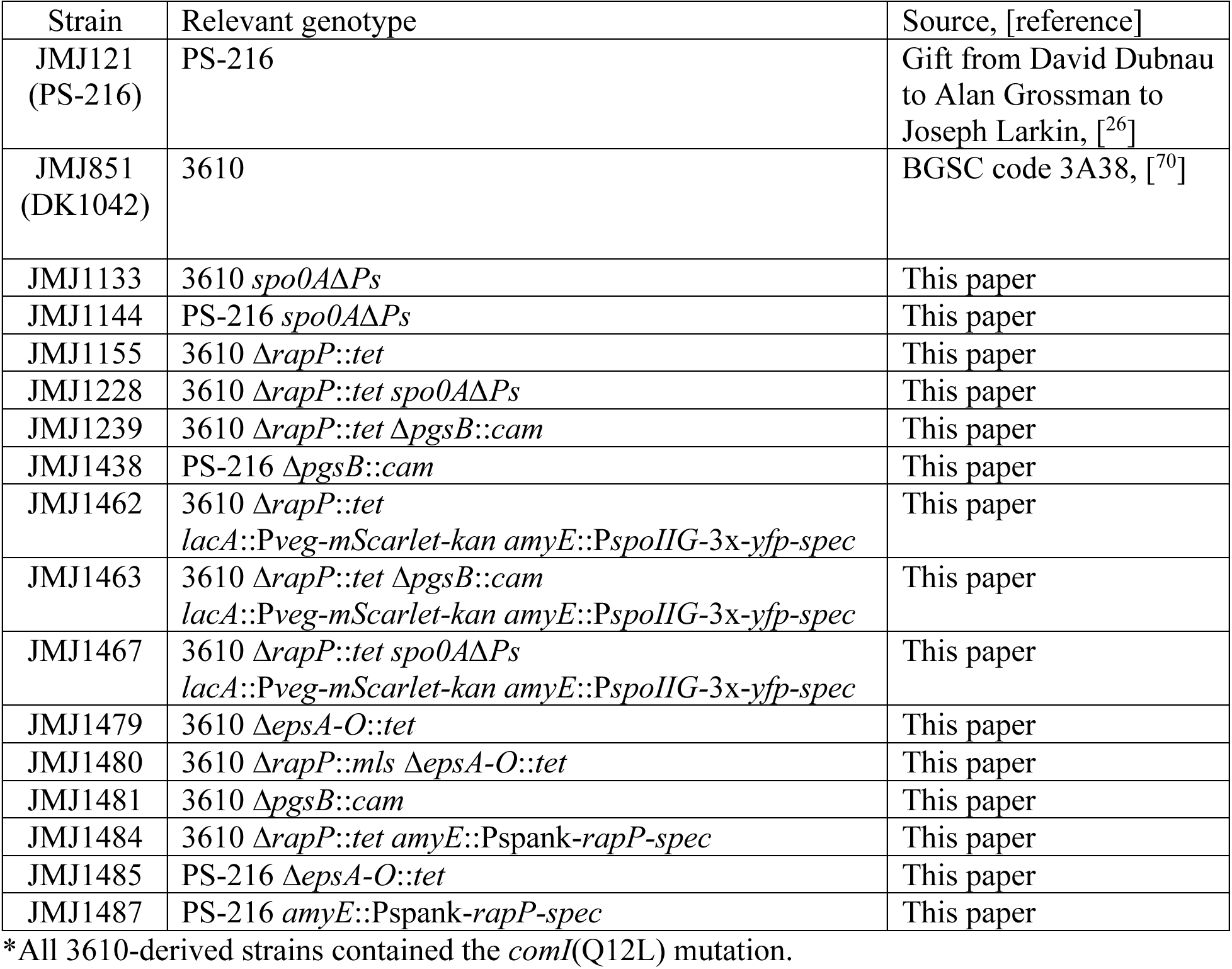
B. subtilis strains used*.

### Construction of knockout mutants

The *rapP*, *pgsB*, and *epsA-O* knockouts were generated by allelic replacement of the indicated genes with antibiotic resistance genes. Homology arms (∼2 kb) upstream and downstream of the indicated open reading frames (ORF) were cloned and fused to an antibiotic resistance gene cassette by isothermal assembly^72^. The resulting DNA fragment was introduced to *B. subtilis* by transformation and antibiotic selection. The *rapP* knockouts eliminated the *rapP* ORF plus 37 bp upstream and 49 bp downstream.

### Construction of inducible *rapP* construct

The *rapP* ORF was cloned into pDR110 (vector for IPTG induction and integration at *amyE*). The resulting plasmid (pJMJ1135) was linearized and introduced to *B. subtilis* by transformation and selection for spectinomycin resistance.

### Construction of constitutive reporter

To label cellular biomass, a strong transcriptional unit composed of the P*veg* promoter and R0 ribosome binding site^73^ was used to drive expression of *mScarlet-I*^74^. The P*veg*-R0-*mScarlet* fragment was fused to a kanamycin resistance cassette and homology arms for integration at *lacA* as previously described.

### CFU plating experiments

Cells were washed from lawns and diluted to an OD600 of 0.5 in 1x Spizizen’s salts. The resulting suspensions were spotted (1 μl) onto the center of the MSgg-agarose substrates and allowed to dry. The well-plates were sealed with BreathEasy Films (Diversified Biotech) and incubated at 30°C. At the time of inoculation, inoculums were diluted and plated in duplicate on LB-agar plates to determine the initial CFU. After the indicated incubation period, biofilms were scraped from the substrate surface and mechanically disrupted with sterile wooden sticks, then resuspended in 0.5 ml 1x Spizizen’s salts. Biofilm suspensions were diluted and plated to determine the total CFU/ml. Spore frequency was determined by plating the same dilutions after a heat treatment at 85°C for 20 minutes to kill vegetative cells.

### Whole biofilm imaging and radial expansion measurements

Biofilms were photographed using an Olympus SZX7 stereomicroscope with a 1x, 0.1 NA objective. Identical zoom was used across sets of samples. Images of a ruler were used to set the pixel-to-μm scale of the images. Samples were illuminated with a ring light mounted above the stage. Images were cropped manually (ImageJ) to remove the substrate surrounding biofilms. For radial expansion measurements, phase contrast images were taken of the dried inoculum spots and mature biofilms using an Olympus IX83 inverted widefield microscope with a 4x, 0.1 NA objective. The area of substrate covered by biomass was measured by manually tracing biofilms (ImageJ) and used to calculate the effective radius of the initial spots and final biofilms.

### Confocal microscopy

Fluorescence images were acquired with a Leica Stellaris 5 confocal system and DMi8 inverted microscope. Biofilms were prepared as described above for CFU plating. For co-cultures, cell density-normalized inoculums were mixed at the indicated ratios prior to spotting on the substrate. Timelapse imaging was started approximately one hour after inoculation, which allowed for warming of the samples to 30°C in the stage-top incubator (Okolab K-frame). The interval between acquisitions was 90 minutes. All fluorescence data (except that used in Figure 6) were acquired with a 25x water-immersion objective. For these experiments, an objective heater was also used. The immersion water was maintained between the objective and the sample plate bottom using silicone tubing (0.64 mm inner diameter, 1.19 mm outer diameter) and a syringe pump with the flow rate adjusted to compensate for evaporation. Regions of interest (ROIs) were defined with sufficient margins to account for the anticipated growth of biofilms as well as the gradual Z-axis drift of samples due to evaporation. Excitation lines for YFP and mScarlet channels were scanned simultaneously, preventing drift between acquisitions. Details of the acquisition settings are listed in Table 2.

**Table 2.**
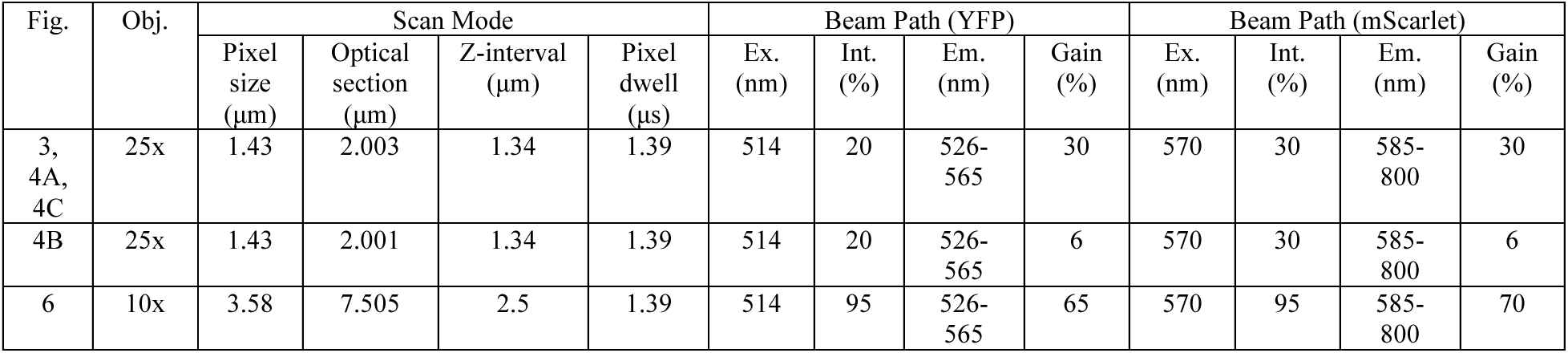
Confocal acquisition settings.

| Fig. | Obj. | Scan Mode |  |  |  | Beam Path (YFP) |  |  |  | Beam Path (mScarlet) |  |  |  |
| --- | --- | --- | --- | --- | --- | --- | --- | --- | --- | --- | --- | --- | --- |
|  |  | Pixel size (μm) | Optical section (μm) | Z-interval (μm) | Pixel dwell (μs) | Ex. (nm) | Int. (%) | Em. (nm) | Gain (%) | Ex. (nm) | Int. (%) | Em. (nm) | Gain (%) |
| 3, 4A, 4C | 25x | 1.43 | 2.003 | 1.34 | 1.39 | 514 | 20 | 526-565 | 30 | 570 | 30 | 585-800 | 30 |
| 4B | 25x | 1.43 | 2.001 | 1.34 | 1.39 | 514 | 20 | 526-565 | 6 | 570 | 30 | 585-800 | 6 |
| 6 | 10x | 3.58 | 7.505 | 2.5 | 1.39 | 514 | 95 | 526-565 | 65 | 570 | 95 | 585-800 | 70 |

### Image processing and analysis

Custom Python scripts were used to subtract background and to generate Z-projections and cross-section slices from confocal z stacks. To distinguish fluorescent biomass from substrate-derived background, which varied over time, we computed thresholds for each timepoint. The brightest background came from the substrate-air interface. We measured this by generating maximum intensity Z-projections, then taking the median and median absolute deviation (MAD) of an ROI positioned ∼20 μm from the biofilm leading edge. The size of this ROI was ∼40 x 40 μm for 25x datasets (Figures 3 and 4) and ∼70 x 100 μm for 10x datasets (Figure 6). Threshold-passing intensities were those that exceeded the substrate-ROI *median* + (*n* × *MAD*). For 25x datasets, *n* was set to 4 for both channels. For 10x datasets, *n* was set to 1 and 2 for YFP and mScarlet channels, respectively.

Z-projections were generated by averaging or summing threshold-passing intensities across z stacks. Radius-Z cross-sections were generated by slicing z stacks along the radius; slices were averaged over their ∼40 μm width (28 pixels for 25x data, 12 pixels for 10x data) to form a 2D image. Numerical radial profiles were derived from the same averaged slices. SciPy functions were used for analysis of radial profiles: smoothing was done with gaussian_filter1d, peak detection was performed with find_peaks. The vegetative cell front was defined from the leading edge of biomass to the “left base” returned by the find_peaks function.

To compare expansion rate and sporulation dynamics in co-culture biofilms, we defined a sporulation front to track the spread of high sporulation activity. The sporulation front was derived from smoothed radial profiles of YFP/mSc and was defined as the most distant (from the biofilm center) position with 75% of the maximum YFP/mSc value. To compare sporulation-front and expansion velocities, boundaries were set to limit the position vs. time data used. The distal boundary was typically defined as the inflection point dividing expansion into two phases (fast and slow). This was done to exclude velocity ratio measurements derived from the slow phase of expansion, which were close to 1 and varied little across samples. The inflection points were defined by finding the most parsimonious two-phase linear description for each leading-edge position vs. time curve. In some datasets (high PGA-producer %), the measured sporulation-front position transiently regressed toward the center due to noise in YFP/mScarlet profiles. In those cases, the boundary was adjusted to the last position before retrograde movement. To smooth instantaneous velocity measurements, the temporal resolution of the position vs. time data was increased 4-fold by linear interpolation (numpy.interp).

### Active fluid model of nutrient-coupled growth and sporulation

Biofilms were represented as expanding discs with fixed height, radial symmetry, and a defined boundary *R*(*t*). Each radius *r* had a fraction *ρ*_1_(*r*, *t*) of cells, a fraction *ρ*_2_(*r*, *t*) = 1 – *ρ*_1_ of spores, and nutrients *n*(*r*, *t*). Cells and spores are confined within *R*, which expands over time due to production of new cells, whereas nutrients are not confined and move freely across the substrate-biofilm boundary. Flux of cells and spores is determined by the velocity field *v*, indicating the effects of newly produced cells pushing other components radially. Flux of nutrients is determined by diffusion *D*. The remaining terms are metabolic reactions that act as either source terms or sink terms for the corresponding dynamical equations. To mimic the experimental geometry, biofilms were initialized with a radius *R*_0_ = 1.14 mm in an environment with radius *L* = 7.8 mm corresponding to experimental growth chambers. The parameters used, except those specified in Figures 5 and 6, were *g* = 1/hr, *n_g_* = 1, *γ* = 4, *n_k_* = 0.03, *h* = 4, *D* = 300 μm^2^/sec. Due to radial symmetry of the model, 1D arrays were sufficient for numerically solving the model. The dynamic variables at each location and time were updated according to a straightforward discretization of the dynamical equations. The code was implemented in Python. Detailed numerical solution steps are included in Supplemental Information.

## Acknowledgments

This work was supported by NIH R35GM142584 (J.WL.), a Burroughs Wellcome Fund CASI award (J.W.L.), NSF PHY-2118561 (A.M.), and NIH R35GM156451 (A.M.). We thank Shila Banerjee, Pankaj Mehta, and members of the Larkin Group for helpful discussions.

## Author Contributions

Conceptualization, J.M.J, M.Y., A.M., J.W.L.

Formal analysis, J.M.J., M.Y.

Funding acquisition, A.M., J.W.L.

Investigation, J.M.J., M.Y.

Methodology, J.M.J.

Project administration, A.M., J.W.L.

Resources, J.M.J.

Supervision, J.W.L.

Visualization, J.M.J., M.Y., A.M., J.W.L.

Writing – original draft, J.M.J., M.Y., A.M., J.W.L.

Writing, review & editing, J.M.J., M.Y., A.M., J.W.L.

---

**Figure S1.**
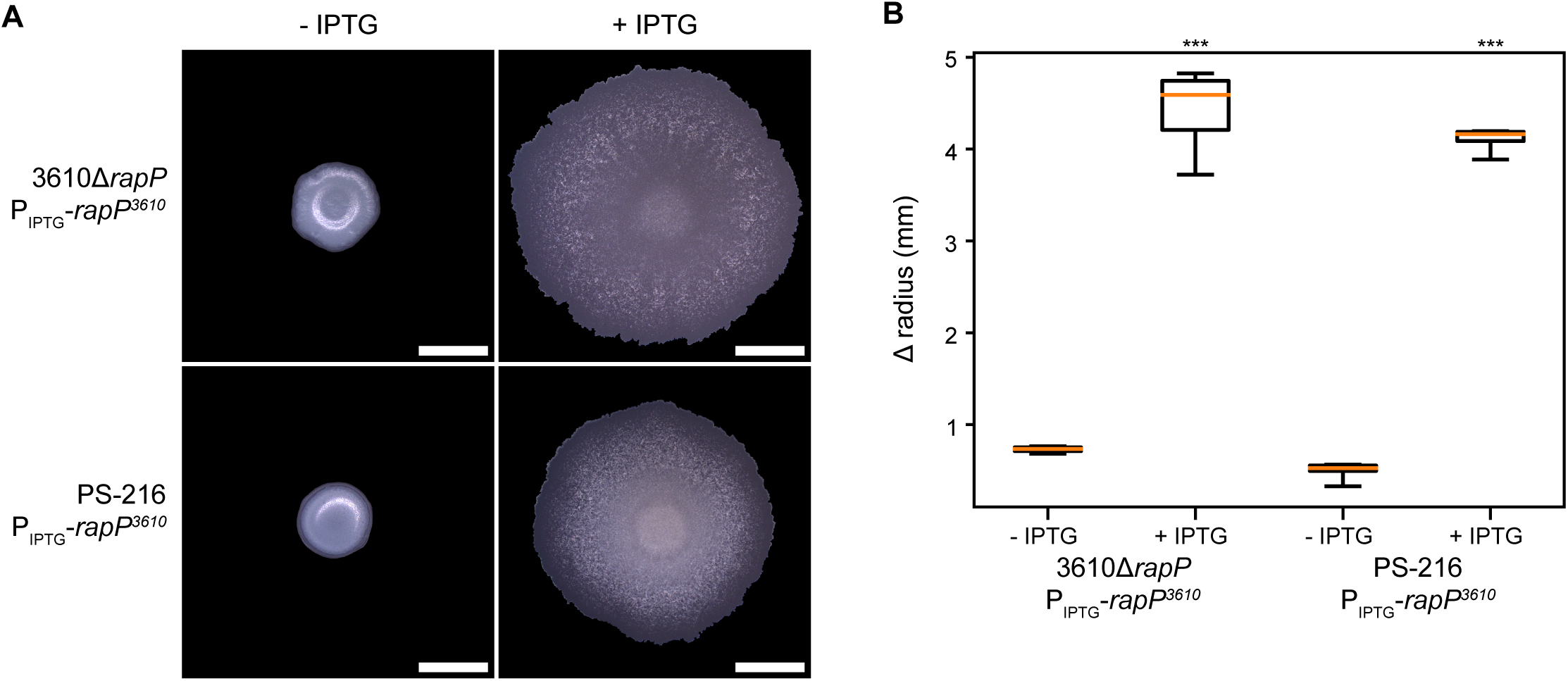
Defective cell-cell signaling alters biofilm morphology. (A) Color photographs of the top surface of biofilms at 24 hpi. (B) Boxplots representing the change in biofilm radius from inoculation to 24 hpi. Central lines indicate medians, boxes enclose the middle 50% of the data, and whiskers indicate the minimum and maximum measurements; *n* = 6 biofilms. *\*\*\*p* < 0.001 by doubled-sided t-test on independent samples with unequal variance (*p* = 4.24 x 10^-6^ for 3610Δ*rapP*+*rapP*, *p* = 3.14 x 10^-13^ for PS-216+*rapP*). Scale bars 3 mm.

**Figure S2.**
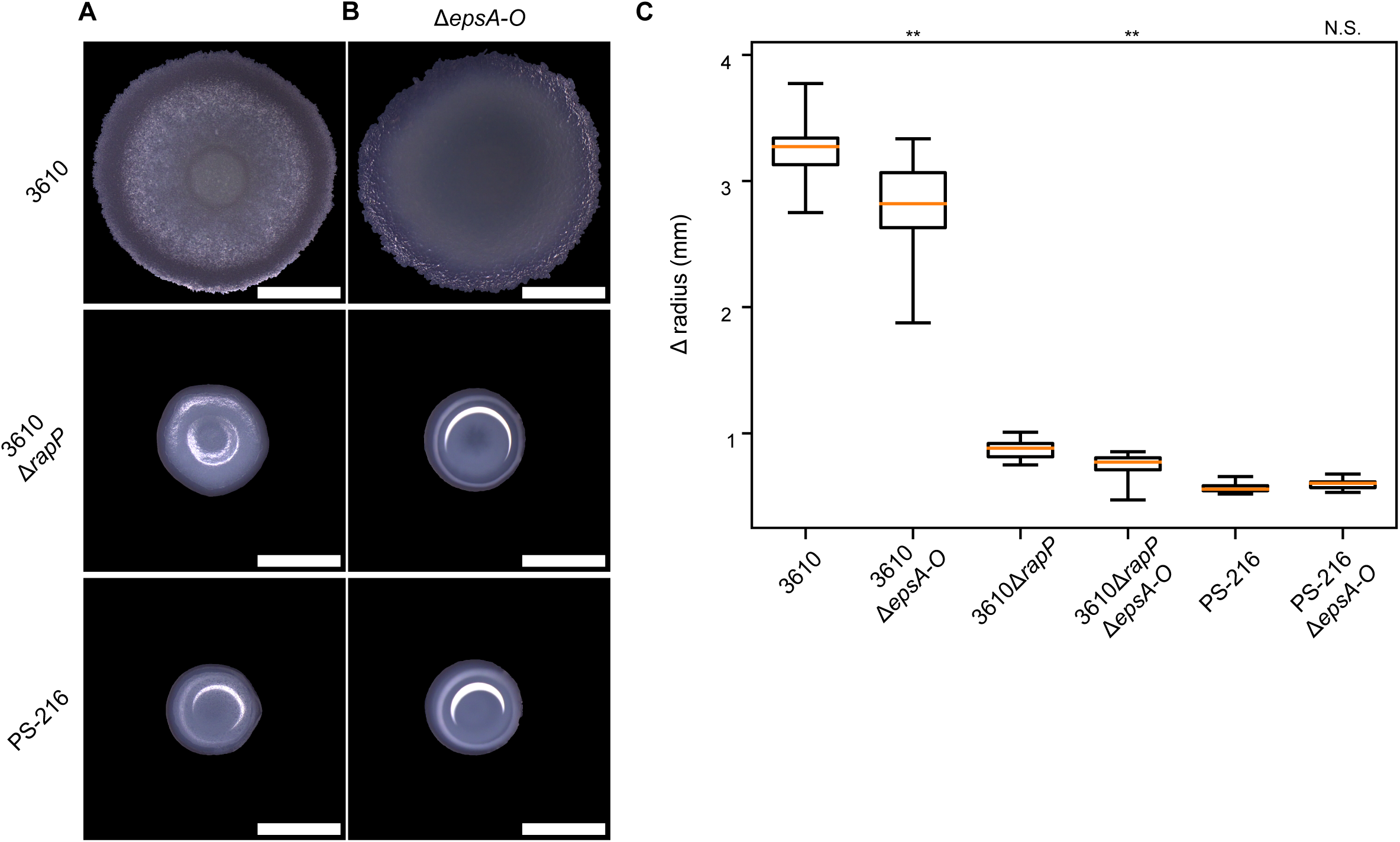
Biofilm spreading is weakly dependent on EPS for strains without *rapP^3610^*. (A-B) Color photographs of the top surface of biofilms at 24 hpi. (C) Boxplots representing the change in biofilm radius from inoculation to 24 hpi. Central lines indicate medians, boxes enclose the middle 50% of the data, and whiskers indicate the minimum and maximum measurements; *n* = 12 biofilms. *\*\*p* < 0.01 by doubled-sided t-test on independent samples with unequal variance (*p* = 0.0049 for 3610Δ*epsA-O*, *p* = 0.0038 for 3610Δ*rapP*Δ*epsA-O*, *p* = 0.101 for PS-216Δ*epsA-O*. Scale bars 3 mm.

**Figure S3.**
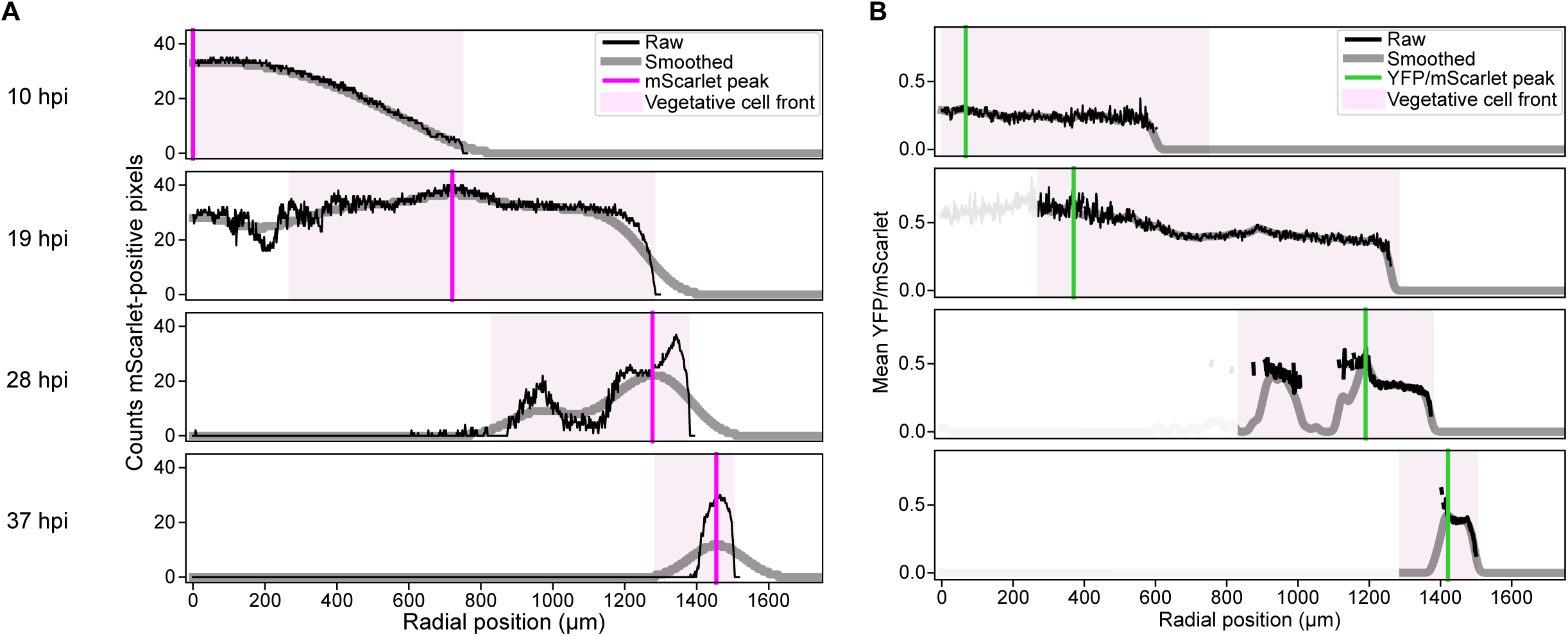
Tracking spread of sporulation activity through biomass. (A) Profiles of mScarlet-positive pixel count vs radial position (related to Figure 4A). One-dimensional gaussian smoothing (*σ* = 50) was used prior to peak detection. Vertical magenta lines indicate the detected peak of the smoothed data. Light magenta shading indicates the vegetative cell front, extending from the left base of the mScarlet peak to the leading edge of biomass. (B) Profiles of mean YFP/mScarlet vs radial position corresponding to the data shown in Figure 4C. Vertical green lines indicate the maximum value of smoothed profiles (*σ* = 6), excluding data outside the vegetative cell front.

**Figure S4.**
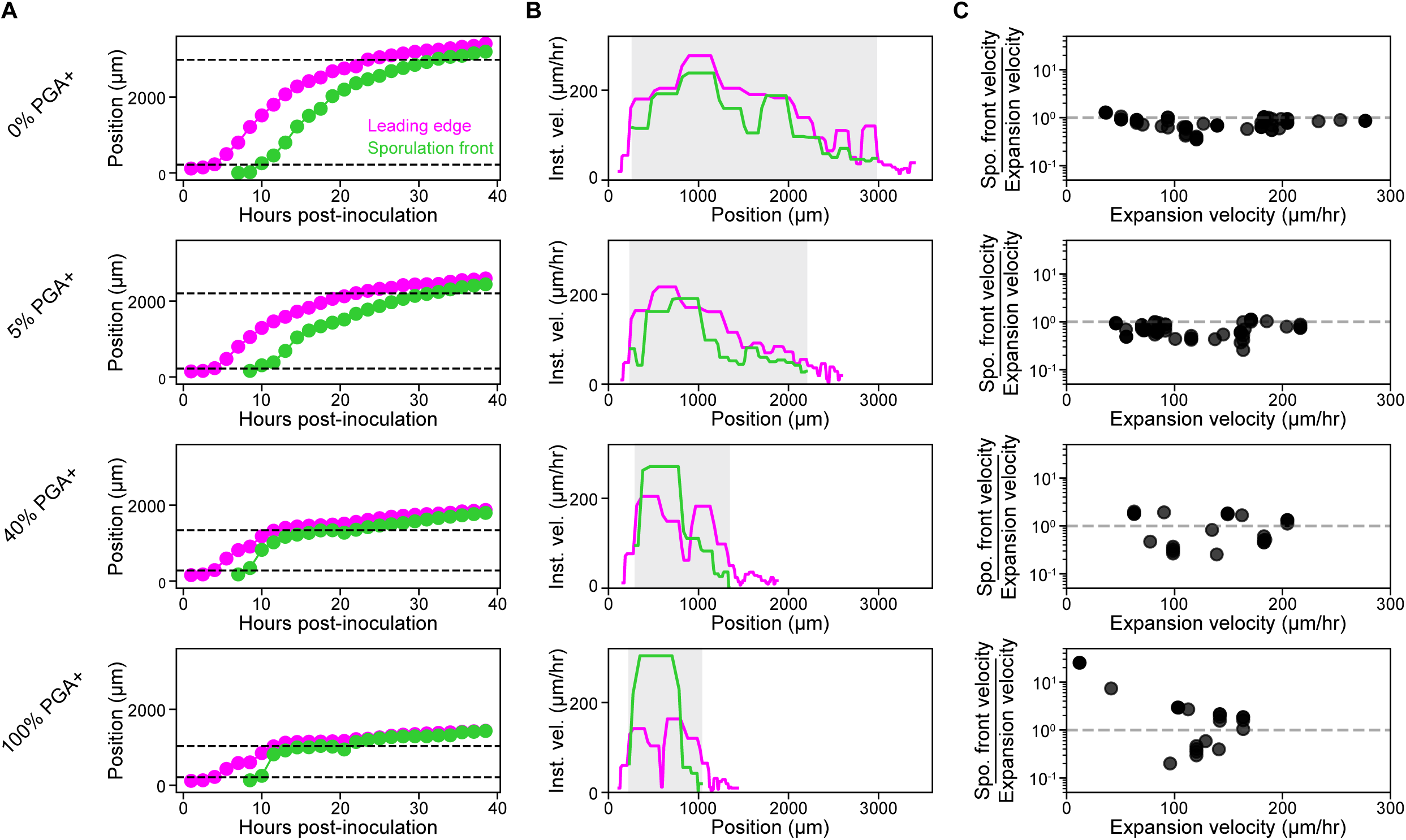
Quantification of leading edge and spore-front velocities. (A) Tracking of biofilm leading edge and sporulation front positions. Magenta and green circles, respectively, indicate experimentally measured biomass leading edge and sporulation front positions. Central lines indicate interpolated values. Horizontal dotted lines mark the range of positions used for calculating velocity ratios. (B) Instantaneous velocity as a function of position for interpolated values in panel A. Gray shading corresponds to the range defined in A by dotted lines. (C) Calculated velocity ratios (sporulation front over expansion) plotted against the corresponding expansion velocities using instantaneous velocity data from the shaded regions in panel B.

**Video S1. Propagating front of mScarlet-positive biomass (related to Figure 4)**

Video running through timelapse frames of mScarlet radius-Z cross-section data. Data are from an independent biofilm compared to that in Figure 4. Images were straightened (ImageJ) to correct for substrate curvature.

Scale bar 100 x 50 μm.

**Video S2. Propagating mScarlet and sporulation activity (related to Figure 4)**

Video running through timelapse frames corresponding to Figures 4A and 4C. Magenta, mScarlet summed across z stacks; green, YFP/mScarlet for dual YFP- and mScarlet-positive pixels.

Scale bar 0.2 mm.

**Video S3. Propagating mScarlet and sporulation activity for single-strain and co-culture biofilms (related to Figure 6)**

Video running through timelapse frames of merged radius-Z cross-sections of mScarlet (magenta) and YFP/mScarlet (green) data for the representative biofilms shown in Figure 6. Images were straightened (ImageJ) to correct for substrate curvature.

Scale bar 500 x 100 μm.

