## Supplemental Information for "Colony morphogenesis regulates sporulation dynamics in bacterial biofilms"

### Supplemental Information for "Colony morphogenesis regulates sporulation dynamics in bacterial biofilm"

#### 1 Detailed Modeling

Biofilms are communities of cells embedded in self-produced extracellular matrix (ECM) polymers. There have been previous studies on properties of cell clusters modeled as active fluids [4, 9–11]. Since we are interested in how *B. Subtilis'* biofilm can be affected by sporulation and ECM production, particularly in terms of biofilm sizes and compositions, we use an active fluid model with a defined boundary to model cells and spores clusters. In addition, we introduce nutrients to represent clusters' growth environment.

##### 1.1 Simplified 2D radial symmetric biofilm model

A biofilm on a Petri-dish grows radially and vertically. Our theory is concerned only with the radial component of growth, and its planar shape is roughly circular during growth. Therefore, we model the biofilm as a radially symmetric expanding circle.

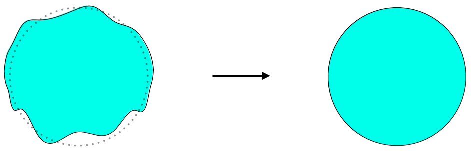

The continuity equation for material inside the biofilm can be written as:

$$\dot{\rho} + \nabla \cdot (\vec{v}\rho) = f(source) + f(sink), \tag{1}$$

where $\dot{\rho}$ is the change of material amount at certain location, $\nabla \cdot (\vec{v}\rho)$ indicates that flux of material through that location, and $f(source)$ and $f(sink)$ represents generation and disappearance of that material.

We have applied this equations to three components – cells, spores, and nutrients in our model. Their interactions are governed by metabolic terms that either act as source terms or a sink terms and physical movements shown later as $\nabla \cdot (\vec{v}\rho)$ (physical pushing of cells and spores) and $D\nabla^2$ (diffusion of nutrients).

#### 1.2 Metabolic terms

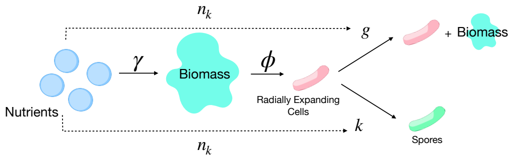


While cells, spores, and nutrients are metabolically associated with each other through complex genetic networks [3, 6, 7], we propose the above simplified metabolic relationship between the three different components.

First of all, cells uses nutrients to make new biomass (including new cells, extracellular matrix (ECM) and other polymers [8]). The efficiency of cells utilizing nutrients is defined by the nutrient conversion rate $\gamma$.

Secondly, because the biggest effect of ECM on biofilm expansion is the inhibition of radial expansion by extracellular poly-$\gamma$-glutamate (PGA), we require in our model that PGA production does not contribute directly to biofilm radial expansion and instead radial expansion is mainly driven by the incompressibility of cells and spores for simplicity. Therefore, we introduce $\phi$ as a parameter that defines how much newly generated biomass is radially expanding cells. The smaller $\phi$ is , the more constrained radial expansion becomes.

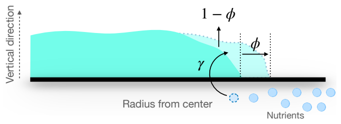


Thirdly, under limited nutrients conditions (implied by $n_k$), proliferating cells can sporulate into spores at a rate $k(n)$. In addition, cells are generating new biomass at a rate $g(n)$.

#### 1.3 Physical interactions and Dynamical equations

Besides cells are proliferating and spores are in a dormant state, we assume that cells and spores have similar physical properties, like density and size, so that they can be pushed by other cells in the same way and are subject to the same velocity field $v$. To simplify their interaction, we demand that they are incompressible. Meanwhile, nutrients do not physically interact with cells or spores, but they diffuse radially.

Therefore, we introduce the following dynamical equations for the 2-D planar expanding biofilm:

$$\dot{\rho_1} + \nabla \cdot (\vec{v}\rho_1) = g\phi \frac{n}{n+n_g}\rho_1 - k\frac{1}{(\frac{n}{n_k})^z + 1}\rho_1 \tag{2}$$

$$\dot{\rho_2} + \nabla \cdot (\vec{v}\rho_2) = k\frac{1}{(\frac{n}{n_k})^z + 1}\rho_1 \tag{3}$$

$$\dot{n} = D\nabla^2 n - g\gamma \frac{n}{n+n_g}\rho_1 \tag{4}$$

$$\begin{aligned}\rho_1 + \rho_2 &= constant \\ &= c \quad within\ biofilm,\end{aligned} \tag{5}$$

where $\rho_1$ represents cells, $\rho_2$ represents spores, and $n$ is the nutrients density. Cells' and spores' physical interaction due to incompressibility is indicated as a generated flow of cells and spore with $\vec{v}$. In addition, new radially expanding cells are generated at an effective growth rate $g\phi\frac{n}{n+n_g}$, and spores are converted from cells at an effective sporulation rate $k\frac{1}{(\frac{n}{n_k})^z+1}$. In addition, $\rho_1$ and $\rho_2$ can also be viewed as scaled fractions to some density when $c = 1$ (which is used in the main text results). On the other hand, nutrients are diffusing with diffusion coefficient $D$ and are used by cells at rate $g\gamma\frac{n}{n+n_g}$.

### 2 Setting of Parameter Values

#### 2.1 Brief summary of parameter values used

| *strain* | $R_0$ | $L$ | $D$ | $g$ | $n_g$ | $\gamma$ | $\phi$ | $k$ | $n_k$ | $h$ |
|---|---|---|---|---|---|---|---|---|---|---|
| 3610 | 1.14mm | 7.8mm | $300\mu m^2/sec$ | 1/hr | 1 | 4 | 1 | 0/hr | 0.03 | 4 |
| $3610\Delta rapP$ | 1.14mm | 7.8mm | $300\mu m^2/sec$ | 1/hr | 1 | 4 | 0.3 | 10/hr | 0.03 | 4 |

#### 2.2 $R_0\&L$: Initial radius of biofilm and simulated space

From the experiments, the initial radius of an inoculum is around $1.1mm$ and the radius of the petri dishes used is around $8mm$. Therefore, in order to use similar values as experiment settings as well as to accommodate the discreteness of simulation, we use $R_0 = 1.14mm$ as the initial radius of the biofilm and $L = 7.8mm$ as the size of the whole simulation space. There are no initial spores present, only cells.

#### 2.3 $D$: nutrient diffusion coefficient

The diffusion coefficient for glucose in water is around $600\mu m^2/sec$ [12]. We assume that the diffusivity of glucose and other similar nutrient sources in agar is smaller than that in water. Therefore, we use $D = 300\mu m^2/sec$ as the diffusion coefficient of nutrients in the simulation.

#### 2.4 $\phi_{3610}$: fraction of biomass used to produce cells – strain 3610 and $3610\Delta rapP\Delta pgsB$

As shown in the model, cells not only use nutrients to produce new cells, but also they can use nutrients to make PGA. Making more PGA corresponds to a lower $\phi$, indicating that a smaller amount of biomass made from nutrients will be cells (that expand the biofilm radially).

In the experiment, the effect of $\phi$ can be observed from strain with different mutations of the *pgsB* gene that is essential for PGA synthesis. For example, strain 3610$\Delta rapP \Delta pgsB$ can form larger biofilms compared to strain 3610$\Delta rapP$, which indicates that 3610$\Delta rapP$ has smaller $\phi$ and produces more PGA than 3610$\Delta rapP \Delta pgsB$. However, the data also shows that strain 3610 and strain 3610$\Delta pgsB$ form biofilms of similar size, despite the fact that 3610$\Delta pgsB$ has the PGA synthesis gene deleted. Therefore, we argue that strain 3610 and strain 3610$\Delta pgsB$ have similar $\phi$. This is consistent with previous observations that strain 3610 represses *pgsB* under standard lab biofilm conditions. For simplicity, we set their $\phi$ value to 1, implying that all biomass produced from nutrients contributes to radial expansion.

Figure 2 - D

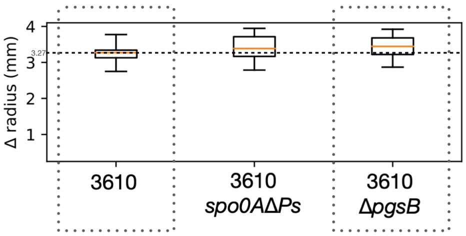


#### 2.5 $\gamma$: Nutrient to biomass/ Nutrient to "radius" conversion rate

To estimate the nutrient conversion rate $\gamma$, we use $\Delta R$ for strain 3610. Since this strain does not sporulate, we assume all available nutrients in the agar plate will be converted into biomass. In addition, since $\phi$ for strain 3610 is 1, all biomass generated will be radially expanding cells and the amount reflects the change of the biofilm size. Thus,

$$\overset{nutrient\ used}{n_0 L^2 \pi} = \overset{cell\ produced}{\frac{\gamma}{\phi} c[(\Delta R + R_{initial})^2 - R^2{}_{\text{initial}}]\pi}, \tag{6}$$

where $n_0$ is the initial nutrient concentration, $L$ is the radius of agar culture plate which is about $8mm$, and $c$ is the local density of cells and spores.

Therefore,

$$\frac{\gamma}{\phi} = \frac{n_0 L^2}{c[(\Delta R + R_{initial})^2 - R^2{}_{\text{initial}}]}. \tag{7}$$

Since the initial nutrient concentration $n_0$ is kept unchanged across experiments, we can arbitrarily set it to 1. For simplicity, we can also scale the cells and spore density $c$ to 1 in regards to $n_0$, since the final size of biofilm of the non-sporulating strain depends only on the ratio $\frac{\gamma c}{\phi n_0}$. In this setting, the value of $\frac{\gamma}{\phi}$ will tune the maximum size of biofilm for a particular strain. Here, considering strain 3610, $\phi$ is set to 1. As a result, we can use the values of $L$, $\Delta R$, and $R_{initial}$ from experiments in the above expression, to find that $\gamma \approx 4$.

Figure 2 - D

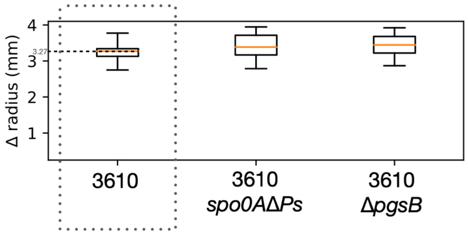


#### 2.6 $\phi_{3610\Delta rapPspo0A\Delta Ps}$: fraction of biomass used for radially expanding cells– strain $3610\Delta rapPspo0A\Delta Ps$

Besides 3610 and $3610\Delta pgsB$, strain $3610\Delta rapPspo0A\Delta Ps$ does not sporulate. Nevertheless, it forms a much smaller biofilm compared to the other two strains, which within our model reflects the effect of PGA production. Therefore, $3610\Delta rapPspo0A\Delta Ps$ has a smaller $\phi$ value in our model. Previously, it has been shown that $\frac{\gamma}{\phi}$ determines the maximum radial growth of biofilm. With $n_0 = 1,\ c = 1$, and $\gamma = 4$ as indicated from strain 3610 and $3610\Delta pgsB$, $\phi_{3610\Delta rapPspo0A\Delta Ps} \approx 0.2$.

However, $\phi_{3610\Delta rapPspo0A\Delta Ps} \approx 0.2$ is gained with the assumption that all nutrients are consumed. In simulation, we find that results produced using $\phi_{3610\Delta rapPspo0A\Delta Ps} \approx 0.3$ reproduces the experimental observations better. This is probably due to the fact that nutrients at the edge of the agar plate take time to diffuse into the biofilm. With the assumption that the nutrient diffusion coefficient is about $300\mu m^2/s$, the characteristic length of diffusion at 60 hours post-inoculation when the data is collected is $\sqrt{Dt} \approx 8mm$, which is about the radius of the agar plate. Thus, it is likely that not all nutrients are used in this case.

Figure 2 - D,E

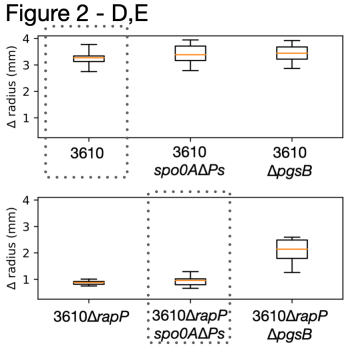

#### 2.7 $\phi_{3610\Delta rapP}$: fraction of biomass used for radially expanding cells – strain $3610\Delta rapP$

From the data, we also notice that strain $3610\Delta rapP$ forms biofilms of similar size as strain $3610\Delta rapPspo0A\Delta Ps$. The difference between the two strains is that $3610\Delta rapP$ can sporulate and $3610\Delta rapPspo0A\Delta Ps$ does not. Nevertheless, the experimental data show that sporulation is not the determining factor for biofilm sizes. The empirical reason is that cells are maintained at the biofilm edge and are exposed to external nutrients. This is consistent with our model, where the maximum biofilm size $(\frac{\gamma c}{\phi n_0})^{-1}$ is independent of the sporulation rate $k$. Therefore, we approximate $\phi_{3610\Delta rapP}$ to be 0.3, the same as for strain $3610\Delta rapPspo0A\Delta Ps$.

Figure 2 - E

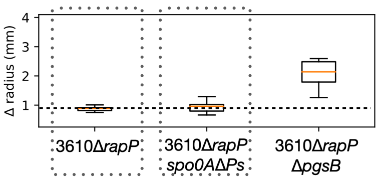


Figure 4 - A

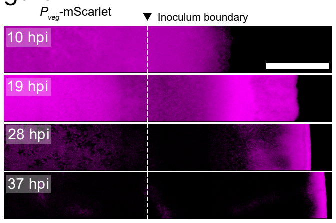


#### 2.8 $k_0$ & $n_k$: sporulation rate

Since sporulation is sensitive to nutrients, and nutrient density is dynamic and radially dependent, it is hard to independently approximate $k_0$ and $n_k$ from experimental data (i.e. the spore percent over time and the sporulation initiation signal profile).

Nevertheless, we do find that $k_0$ and $n_k$ are related to each other and can be estimated together. More specifically, a larger $k_0$ corresponds to a faster depletion rate of the cell signal $\rho_1$ when the nutrient density hits the critical value $n_k$, leading to faster shrinking fronts of cell signal. On the other hand, $n_k$ needs to be small when $k_0$ is large so that the spore percent is small during early times and only goes up quickly at later times. In addition, a large hill coefficient $h$ is needed to keep the sporulation rate near 0 at early times of biofilm expansion when nutrients are abundant.

We tested several sets of $k_0$ and $n_k$ and compared to the cell signal profile at different times and spore percent data from experiments. We find that $k_0 = 10/hr$, $n_k = 0.03$ and $h = 4$ produce qualitatively similar results to those of the experiments.

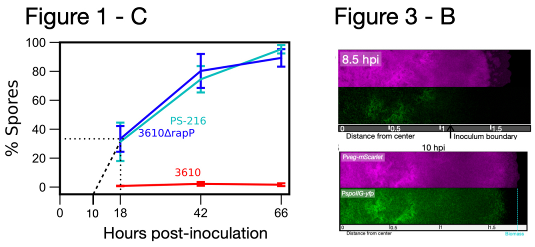


#### 2.9 $g_0$ & $n_g$:growth rate

Like the sporulation rate, it is difficult to independently infer the exact values of $g_0$ and $n_g$ for the model from the experiments. Nevertheless, we assume the reasonable maximum growth rate of $g_0 = 1/hr$ and set $n_g = 1$ to express the fact that the growth rate of *B. subtilis* is sensitive to the nutrient concentration up to and including the initial amount [1, 2, 5].

### 3 Simulation set up / Numerical solution

Since the model is 2D radially symmetric, the profiles of cells, spores, and nutrient on the petri dish only depend on radius $r$. This also applies to any flow of materials. As a result, 1D arrays are sufficient for representing the 2D biofilm expansion while accounting for the choice of using radial coordinates.

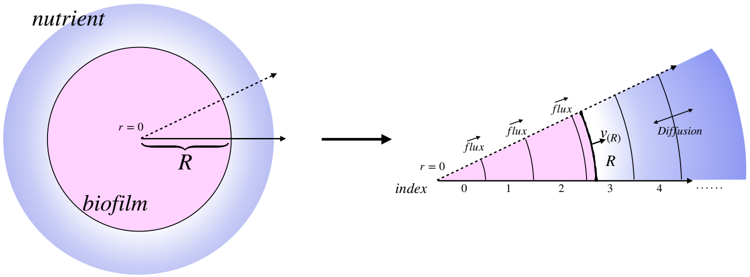


In addition, we assume that cells and spores are incompressible and interact with each other mechanically through pushing, and they are not excluded from the biofilm during expansion (i.e. they do not die). Nutrients are small enough to freely diffuse along the radial direction.

Due to the incompressibility and symmetry, the following rules apply for the simulation:

- For any time point $t$ during simulation when the biofilm edge is at radius $R$, cell profile $\rho_1$ and spore profile $\rho_2$ are valid within $R$. Nutrients density profile $n$ is valid over the whole simulated space.
- Due to incompressibility, flux is induced by production of new cells. In addition, flux can only flow from the center to the edge due to biofilm

symmetry. Therefore, $v$ at a certain location is related to the total flux from $r = 0$ up to the location.

- Cells and spores are pushed in the same way through the local velocity field.

- $R_{new}$ is determined by expansion speed at the edge of the biofilm $v_{(R)}$. $v_{(R)}$ is related to the production of new cells in the whole biofilm.

- There are fluxes of cells and spores within $R_{new}$. There are no fluxes of cells or spores outside the predicted $R_{new}$, because already existing cells and spores with newly produced cells and spores are contained within $R_{new}$ and do not diffuse. However, due to the discreteness of the simulation, flux is restrained within the largest, closest, discretized value of $R$ to $R_{new}$.

- Nutrients are freely diffusing in all simulated space and being consumed by cells, $\rho_1$.

The following diagrams show the update procedure for $\rho_1$, $\rho_2$, and $n$ with imposed rules and conversion relationships.

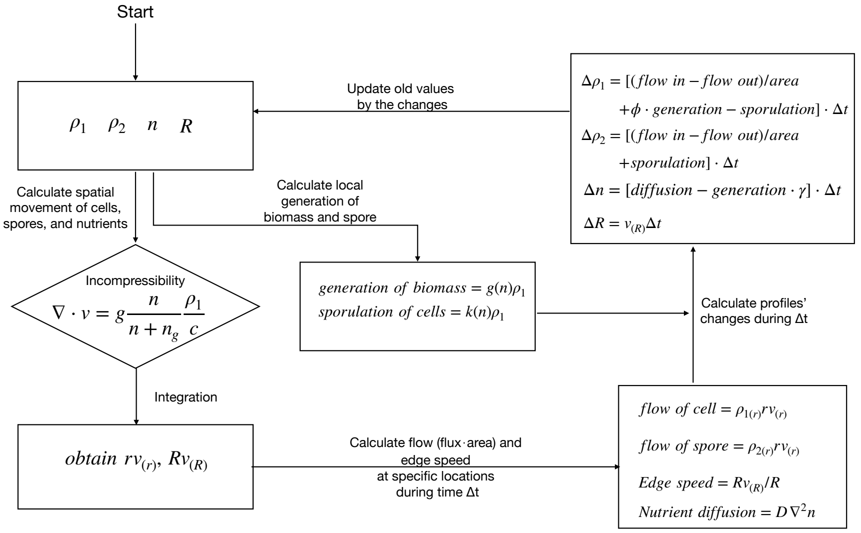


In addition, while the radius is discretized into 1D arrays where the spacing between two nearest indexes is held fixed, the corresponding 2D area of each index is different. This also affects the nutrients available to the biofilm, particularly at the edge. Therefore, making the radius with finer spacing for numerical solution can be crucial for accuracy because $v$, especially $v_{(R)}$, is sensitive to nutrient availability. As shown in the graph below, available nutrients are different for different radial spacing.

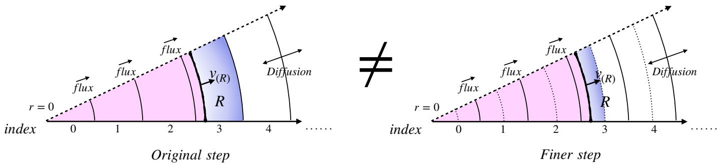


#### 3.1 Special processing used for figure panels

##### 3.1.1 Figure 5C

Spore percent is calculated from $\frac{\sum \rho_2}{\sum(\rho_1+\rho_2)}$.

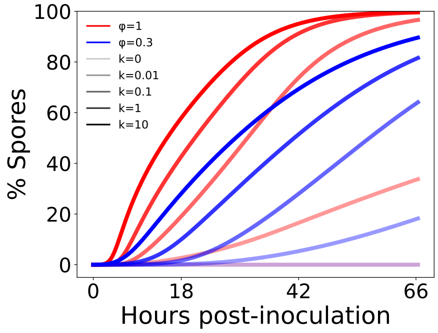


##### 3.1.2 Figure 5D

While the simulation is in the 1D radial direction, the 2D biofilm plot is made by mapping radius from the 1D simulation arrays of cells and spores to corresponding locations in the 2D plane. The gray represents the biofilm area within the biofilm edge defined by the total fraction of cells and spores being greater than 0.01.

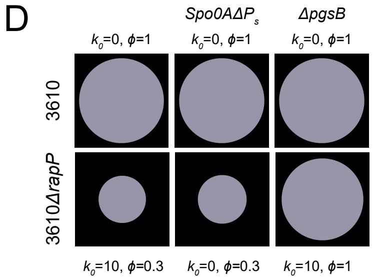


##### 3.1.3 Figure 5E & 5F

The profile plotted is part of the mapped 2D biofilm with data from the 1D simulation results (using a similar technique as Figure 5D). The brightness of color represents the normalized signal value to the max signal value at each time.

The magenta color represents normalized cell signal and green color represents normalized sporulation signal. By using normalized values, the signal is more visible for further analysis.

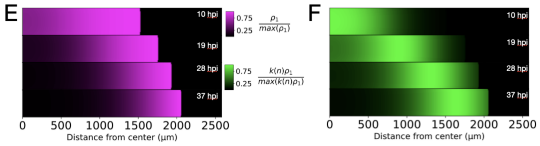


##### 3.1.4 Figure 5G

The data plotted is radius over time for biofilm and sporulation signals. More specifically, for the biofilm, the recorded radius is the furthest position where $\rho_1 + \rho_2 > 0$ from the center $r = 0$. And the radius for the sporulation signal is determined to be the furthest position where the normalized sporulation signal is greater than 0.75 of the maximum sporulation signal at that time across the whole biofilm. To make the graph more concise, we plot the change of radius (i.e. $R - R_0$) starting from 0.5 hours post-inoculation and shade the region where $\rho_1 >= 0.5$.

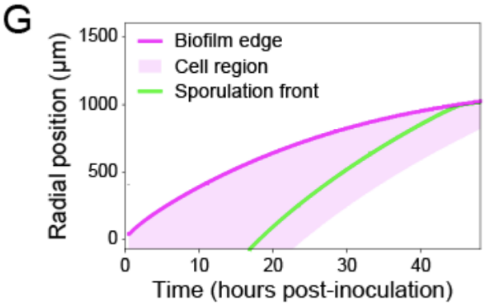


##### 3.1.5 Figure 5H: velocity over time

Due to discretization of radius in the numerical simulation, the average speed of expansion during a certain time frame $\Delta t$ is also discretized. Therefore, in order to plot smoothed velocity of expansion over time, we need to obtain smoothed radius over time. First of all, we only keep the radius data that are greater than $R_0$, the initial radius of the inoculum, so that the $R$ represents the expansion away from the initial inoculum. Afterwards, we obtain a set of times corresponding to $R$ reaching each discretized radial position. The choice of time is when $R$ is about to change to the next value, in which case the actual simulated $R$ is close to the recorded discretized value.

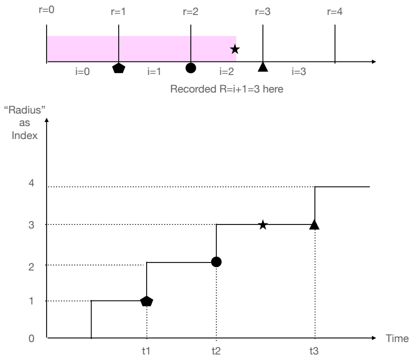


Then, we make a smoother graph of the paired radius and times using B-spline functions with degree of 2 (function/package available in python as "*scipy.interpolate.make_interp_spline*"). Next, we calculate the average velocity $v(t) = \frac{R(t+\Delta t/2)-R(t-\Delta t/2)}{\Delta t}$, where $R$ here refers to the smoothed radius. Nevertheless, we enforce that $v(t=0)=0$ and the calculation for $v(t>0)$ is valid only when $R(t-\Delta t)$ is valid.

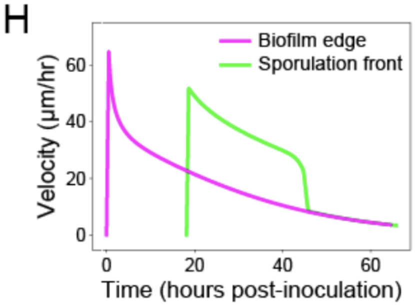


##### 3.1.6 Figure 6A

We plotted the mean average velocity in $\Delta t = 1hr$ time frame for different $\phi$ with black lines representing the standard deviation of velocity collected for each $\phi$. The velocity is calculated from $v(\Delta R, t) = \frac{\Delta R(t+\Delta t/2)-\Delta R(t-\Delta t/2)}{\Delta t}$. Here, $R$ is the recorded discretized radius of either the edge of biofilm or the sporulation signal front. In addition, the pair of $(R,t)$ we choose to calculate speed is where $R$ is about to move to the next discretized radial position. In this case, the actual simulated radius is close to the recorded discretized value.

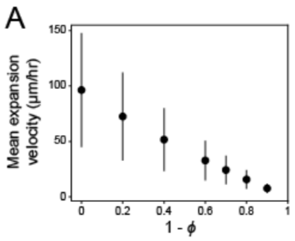


##### 3.1.7 Figure 6B

We've plotted the change of radius of different $\phi$ values. More specifically, we plot $\Delta R$ from the initial radius of the inoculum of the edge of biofilms (defined by $\rho_1 + \rho_2 > 0$) and the front of the sporulation signal (defined by the furthest location in the biofilm where $\frac{k(n)\rho_1}{max(k(n)\rho_1)} = 0.75$). In addition, $\Delta R = R - R_0$ if $R > R_0$ and $\Delta R = 0$ if $R <= R_0$. Here, $R$ is discretized.

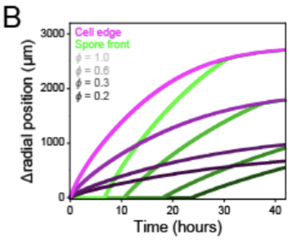


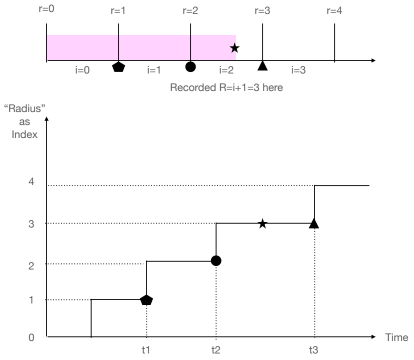


##### 3.1.8 Figure 6C

The $\frac{spore\ velocity}{cell\ velocity}$ is calculated from average biofilm expansion velocities (i.e. cells velocities) and spore signal velocities at the same $\Delta R$ with the same $\phi$ within a given time frame of $1hr$. Here, the $\Delta R$ used for calculation is discretized (unsmoothed). The value then then plotted against the corresponding average biofilm expansion velocities (i.e. cells velocities). Currently, the graph contains the velocity ratios for $\phi = [1, 0.8, 0.6, 0.4, 0.3, 0.2, 0.1]$.

C
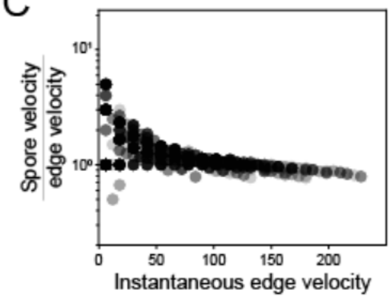
